# Delayed effects of water-limiting conditions influence the ramet demography of a native iterocarpic thistle

**DOI:** 10.1101/2025.05.27.656337

**Authors:** John N. Mensah, Svata M. Louda, Kathleen H. Keeler, Brigitte Tenhumberg

## Abstract

1. Increasing temperatures and shifting precipitation patterns are major components of climate change. Yet, the demographic responses of plants to such changes remain poorly known. We used 20 years of demographic monitoring data (1990–2009) on native *Cirsium undulatum* (wavyleaf thistle) from two Sandhills prairie sites in the central Great Plains, USA, to assess long- and short-term effects of weather variation on ramet dynamics.
2. *C. undulatum* is a deeply taprooted, short-lived perennial plant that can vegetatively reproduce rosette-like ramets. Since field evidence over the full length of the 20 years documented the recruitment and fate of rosettes, we evaluated ramet vital rates and dynamics. We first estimated annual recruitment, survival, and between-stage transitions. These vital rates were used to develop a matrix population model to calculate annual asymptotic population growth rates (λ_t_). We then fitted a functional linear model to explore the effect of temperature, precipitation, and standardized precipitation evapotranspiration index (SPEI), an integrated measure of drought intensity, on ramet demographic rates. Finally, we developed a population viability analysis (PVA) model to evaluate the persistence of the ramet population under worst- and best-case scenarios of future drought events.
3. The results revealed that at the drier site, Arapaho, wetter than normal years over the previous 19 months increased five parameters and decreased one. Seedling recruitment, flowering single rosette sprouts and λ_t_ increased with positive SPEI; and single rosette survival and multiple rosette stasis increased with precipitation. Retrogression from multiple rosette to single rosette was reduced when precipitation decreased. Additionally, higher-than-normal temperatures between 18–12-months period before the population census significantly increased the flowering probability of single rosettes. At the second site, Niobrara, only seedling recruitment was significantly increased by positive SPEI. The PVA indicated *C. undulatum* ramet populations will likely persist even with an increase in drought frequency.
4. *Synthesis:* This study provides a relatively rare long-term study of the effect of weather variables on demographic rates and plant persistence. Our simulations suggest that plants evolved in highly variable and droughty environments like *C. undulatum* will be resilient to increases in drought with climate change.

## 1.0 Introduction

Global climate change is characterized by increasing temperatures and shifting precipitation patterns (Intergovernmental Panel On Climate Change (Ipcc), 2023). Since 1986, the Great Plains region of central North America has experienced temperature increases of 0.42°C - 0.94°C from south to north (Ojima et al., 2021). By 2050, temperatures in this region are projected to rise by more than 2°C, potentially intensifying drought conditions (Ojima et al., 2021). Such climatic changes can profoundly affect plant population dynamics (Doak & Morris, 2010; Frei et al., 2014; Iler & Inouye, 2013); yet, relatively few long-term studies have explicitly examined how demographic rates respond to fluctuations in temperature and precipitation (Crone et al., 2011). Consequently, a deeper understanding of how plant populations respond to climate variability is urgently needed (Tredennick et al., 2021).

Further, the demographic effects of drought and elevated temperatures may be delayed, taking months or years to manifest because plants may adjust their resource allocation and physiological processes over multiple growing seasons (Watts & Tenhumberg, 2021). For example, in dry years plants tend to reduce stomatal conductance to mitigate water loss, a strategy that conserves water but also decreases photosynthetic capacity (Chaves et al., 2003), slowing resource accumulation and potentially delaying demographic processes. Alternately, when conditions shift to a wetter period, plants may efficiently accumulate and reallocate stored resources to growth and reproduction, ultimately influencing long-term population growth rates (Dahlgren et al., 2016). Such delayed effects are common in arid and semi-arid environments; so, arid system plants often require multiple growing seasons to accumulate the resources needed for growth and reproduction (Evers et al., 2021; Tenhumberg et al., 2018). Identifying and understanding the consequences of delayed effects of weather variations is essential for devising appropriate management strategies.

Yet, few studies have examined delays in the effects of weather variables on plant demography (Evers et al., 2021), partly due to the scarcity of long-term data sets that capture interannual variation (Salguero-Gómez et al., 2015). Here, we contribute to this limited body of work by analyzing 20 years (1990 – 2009) of demographic data on *Cirsium undulatum* Spreng. (wavyleaf thistle), a taprooted, iterocarpic native plant in the Sandhills prairie of Nebraska, central Great Plains, USA.

The Nebraska Sandhills, at 50,000 km^2^, form the largest continental dune grassland in the Western Hemisphere, and offer an ideal setting to investigate plant demographic responses to variations in weather variables (Poděbradská et al., 2019; Stephenson et al., 2019). This semi-arid sand prairie has coarse, undeveloped, granitic soils with low water-holding capacity and high susceptibility to wind erosion (Menard et al., 2018). Across the Nebraska Sandhills, the eastern half receives moisture from southerly winds coming across the Gulf of Mexico, whereas the western half is drier and less humid (Shulski et al., 2013).

In recent decades, Nebraska’s climate has shifted toward higher temperatures and more frequent and extreme precipitation events; however, drought remains the predominant abiotic stress (Frankson et al., 2022). This study explored the delayed effects of key weather parameters (precipitation, temperature, and drought) on the demography of *C. undulatum*. We focus on ramet demography because the field observations tracked rosette ramets within plots over two decades; genetic information required to distinguish ramets of different plants (genets), was limited to two years (Brozek, 2009).

We hypothesized that differences in weather conditions between the populations in the northcentral region (Niobrara Valley Preserve) and southwestern region (Arapaho Prairie Preserve) of the Sandhills influenced local-scale variations in ramet demography and their long-term response to weather variation. To test this hypothesis, we estimated demographic vital rates of *C. undulatum* ramets and developed matrix population models (MPMs) to synthesize the effect of individual vital rates on annual asymptotic population growth rates.

The three main questions were: (1) What are the site-related ramet demography and population growth rates? (2) Do temperature, precipitation, and drought have delayed effects on vital rates and population growth rates at either site? and (3) What is the likely long-term effect of variation in drought frequency on *C. undulatum* ramet population viability?

## 2.0 Methods

### 2.1 Natural history

In the central Great Plains, *Cirsium undulatum* Spreng. (wavyleaf thistle, Asteraceae) is an iterocarpic perennial plant with deep taproots (Kaul et al., 2011). It reproduces sexually by seed and asexually (sprouts) by the growth of upper axillary buds on the taproot. Ramets of adjacent taprooted genets intermix in the field, so distinguishing genets is difficult; the required DNA sampling was only done in two years (Brozek, 2009). Thus, in order to take advantage of the long-term field data, we focused on modeling ramet dynamics.

Sexual reproduction occurs via ramet flowering, seed set, and subsequent seedling establishment (Louda, 1998). Flowering begins in June and peaks in July. Seed set occurs in June - July and dispersal occurs in late July - early August. Seedlings emerge the following spring and establish as small rosettes (ramets) in their first year. Unlike invasive weedy thistles, there is no seed bank for the native *Cirsium* spp. at Arapaho Prairie Preserve (Potvin, 1988). A flowering (bolting) ramet is monocarpic and dies after seed maturation, but the taproot can remain alive. Seedlings are distinguished from small asexual sprout ramets by size and the presence of cotyledon leaves.

Asexual ramet reproduction, via sprouts from the underground taproot meristems, forms new ramets on the soil surface in spring (*S. M. Louda, pers. observation*). Most new ramets recruited as sprouts emerged as a single rosette; however, if the apical meristem was damaged early, then the new ramet could be a multiple rosette, composed of two or more sub-rosettes, within 1 cm of each other and with synchronized development. Multiple rosette ramets become single rosette ramets upon the death of all but one sub-rosette.

Wavyleaf thistle ramets all die back each winter. Survivors resprout the following spring, observed at their tag. When a ramet failed to resprout during one growing season but re-emerged at the plant tag within two growing seasons, we treated it as alive but inactive below-ground during that interval (*see* Eriksson, 1988).

### 2.2 Sandhills Prairie Ecosystem and Study Sites

The data on *C. undulatum* ramet dynamics were collected at two native grassland reserves in the Nebraska Sandhills, central Great Plains, Nebraska, U.S.A. The Sandhills is 50,000 km^2^, the largest continental dune grassland in the Western Hemisphere (Muhs & Budahn, 2019), and is composed of eolian sands (92–97% sand soil) deposited thousands of years ago (Miao et al., 2007). Annual precipitation ranges from 450 mm to 690 mm, west to east; however, much of the moisture is lost via evapotranspiration (Shrestha et al., 2021). The sand soils enable high infiltration rates, with annual groundwater recharge rates varying from 40 ± 85 mm in the west to a high of 200 ± 85 mm in the southeast (Wang et al., 2009).

One study site was Arapaho Prairie Preserve (henceforth, Arapaho) in Arthur County, southwestern Nebraska (41°29.733’N, 101°52.502’W). The *C. undulatum* plots were in the valley grassland vegetation between the dunes (Keeler et al., 1980), with fine sandy soils characterized by relatively low infiltration rates. Management at Arapaho involved rotational haying every four years in late July (after the population censuses): 1989, 1993, 1997, 2001, and 2005.

The second study site was Niobrara Valley Preserve (henceforth, Niobrara) in Brown County, northcentral NE (42°43.755’ N, 100°03.754’ W), 270 km northeast of Arapaho. The *C. undulatum* plots were in Sandhills prairie vegetation, with fine sandy soils. The Nature Conservancy’s management of this site involved a sustainable annual grazing regime, involving early-season grazing (4 - 6 weeks: May – mid-June) by a relatively low number of cow-calf pairs (<150 pairs) plus movement of cattle through the site in the fall (Rand et al., 2020, *A. Steuter, personal communication*).

### 2.3 Field data collection

Seven 12 m × 12 m demography plots (144 m^2^ each) were established in 1990: 4 plots at Arapaho and 3 plots at Niobrara. Each wavyleaf thistle ramet in each plot was given a unique numbered tag in 1990; subsequently, when new ramets were encountered, they were individually tagged and measured. For 10 years (1990 - 1999), all ramets were measured twice a year, early growing season (late May) and late season (mid-July). Then, for the next 10 years (2000 – 2009), all ramets were measured once yearly: in late May 2000 - 2004, with only flowering ramets remeasured in July; in mid-July 2005 – 2007; and in late-June 2008 – 2009 at the end of the study; when new ramets no longer received a numbered tag.

For each ramet observed in each census, we recorded stage, size as root crown diameter (mm, using calipers), number of green leaves, length of longest leaf (cm), and plant condition. When a ramet was observed > 10 cm from the nearest existing tag, it was conservatively categorized as a new ramet “sprout” and given a unique number tag.

The demographic stages of the ramets were: seedling, single rosette (SR), multiple rosette (MR), flowering single rosette (SF), flowering multiple rosette (MF), and inactive (IA) (Figure 1). The multiple rosette stage includes ramets with up to five sub-rosettes; however, given the sample sizes available, we did not examine differences in transitions among the multiple rosette ramet types. For analysis, we looked back and categorized a ramet as inactive rather than dead if it reappeared at its tag after being missing 1 or 2 previous growing seasons. If we did not find a ramet for 3 consecutive growing seasons, we assumed it had died when it was first missing. For flowering ramets (bolters), we also recorded stem length (cm) and the number of heads that reached anthesis (flowered). Ramets die after flowering (monocarpic).

**FIGURE 1.**
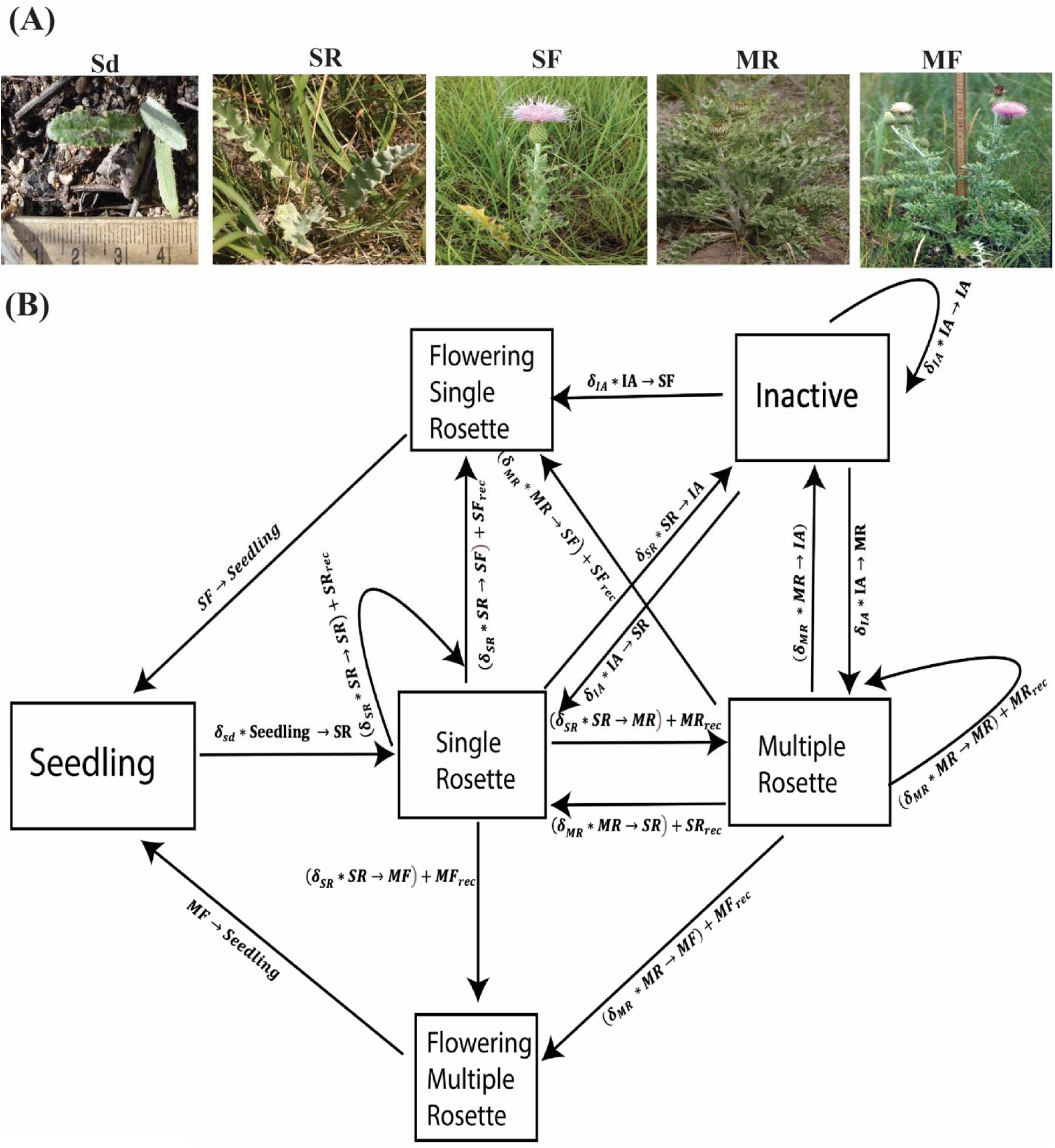
(A) Above-ground demographic stages of *C. undulatum* ramets: Sd = seedling; SR = juvenile single rosette, SF = flowering single rosette, MR = juvenile multiple rosette, MF = flowering multiple rosette; and, IA = inactive stage; (B) Life cycle diagram representing the observed demography of *C. undulatum* ramet stages. Additional symbols represent: for recruits: recruited as a seedling = Seedling_rec,_ and, recruited as a vegetative sprout in stage: MR = MR_rec_, MF = MF_rec_, SR = SR_rec_, and SF = SF_rec_; and, for survivorship: δ_sd_ = seedling survival, δ_SR_ = SR survival, δ_MR_ = MR survival, δ_IA_ = IA survival. Arrows indicate transitions recorded among ramet stages.

We focused our analysis on the demographic stage transitions of the numbered ramets (Details of data management and processing: Appendix S1).

### 2.4 Analyses of demographic data

Data analyses were done with R (R Core Team, 2024). We used exploratory analyses of seedling and sprout recruitment and ramet demography to gain an overview of stage frequencies, i.e., the number of ramets found in each demographic stage each year at each site. We also determined if recruitment, population numbers, and stage frequencies increased, decreased, or remained stable over the study period at each site.

#### 2.4.1 Demographic transitions

We estimated annual transitions between the life history stages late in the *C. undulatum* growing season, specifically July in year *t* to July in year *t*+1. We used Generalized Linear Mixed Models (GLMMs) with year as a random effect (“glmer” function in lme4 package) to estimate annual vital rates for all ramet stages, including recruitment of seedlings and vegetative sprouts (Tenhumberg et al., 2018).

To estimate annual survival probability and stage transitions, we used a binomial error distribution with logit-link. In order to ensure that all possible transition probabilities did not exceed 1.0, we calculated conditional transition probabilities (Statistical details and associated equations: Appendix S1).

To estimate annual recruitment probability, we used a Poisson error distribution with log-link. New vegetative sprout ramets could first appear as a single rosette (SR_rec_), multiple rosette (MR_rec_), flowering single rosette (SF_rec_), or flowering multiple rosette (MF_rec_). These sprout recruits originated from an underground taproot, likely linked to an unknown prior ramet.

We assumed that all seedlings in year *t+1* germinated from seeds produced by flowering ramets ***(f)*** in year *t,* since no *C. undulatum* seed bank occurred (Potvin, 1988). In the few cases where a transition was missed in a particular year, we substituted the estimated overall mean (the model fixed effect) for the missing annual estimate.

We constructed annual matrix population models (MPM) to calculate the asymptotic population growth rates (λ_t_) using all observed annual transitions of the *C. undulatum* life cycle (Figure 1). We also compared the estimated asymptotic population growth rates to the transient growth rates, calculated as the 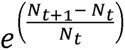, here *N_t_* = ramet population in the current year and *N_t_*_+1_= ramet population in the next year. The transient population growth rate captures short-term changes in the population growth rate. The asymptotic population growth rate assumes that the stable stage distribution (SSD) is reached. If environmental variation reduces or increases each stage by the same percentage the SSD remains the same, and there will be no deviation from the asymptotic population growth rate (Tenhumberg, 2010). (Details on vital rate estimation and development of MPMs in Appendix S1).

### 2.5 Weather data

We used monthly total rainfall and mean monthly temperature data from the weather station at each site, downloaded from the PRISM Climate Group (Daly & Bryant, 2013; https://prism.oregonstate.edu/explorer/). We also calculated the Standardized Precipitation Evapotranspiration Index (SPEI) (Vicente-Serrano et al., 2010). SPEI is a multi-scalar drought index used to assess the onset, duration, and severity of drought events for a selected time scale compared to a long-term average (reference period) in a geographic region (Vicente-Serrano et al., 2010). SPEI captures moisture conditions that deviate from the long-term average, both wetter-than-normal conditions (positive SPEI) and drier-than-normal conditions (negative SPEI).

We used 40 years prior to the study (1950 - 1989) as the reference period and one month as the time scale, the length of the aggregation period over which the total surface water balance was calculated. To get SPEI, we first calculated the potential evapotranspiration (PET) (Thornthwaite, 1948) based on monthly mean temperatures. We then calculated monthly SPEI by quantifying the difference between monthly precipitation (PPT) and monthly PET (Vicente-Serrano et al., 2010) (Details: Appendix S1).

### 2.6 Effect of weather variables on plant demographic parameters

We used functional linear models (FLMs) to identify the relevant time lag—that is, the interval between a climatic event (e.g., drought) and the resultant plant demographic response (Teller et al., 2016). We fitted FLMs (Equation 2) as smooth functions of discrete lagged months to assess the effect of past monthly variation in each of the three weather variables on demographic parameters of *C. undulatum* ramet populations at each site (Teller et al., 2016; Tenhumberg et al., 2018). All FLMs were fitted using the Generalized Additive Model (GAM) (mgcv package, R software) (Details of model fitting and associated codes: Appendix S1).

An FLM fits a smooth spline to the discrete lagged monthly variable *s*(*m*), where *m* is the number of discrete lagged months, in our case beginning from June (*m* = 0; one month before the census month, July) of the monitoring year *t* to a maximum of *L* months in the past. So, the performance of the response variable, *y*, at any year, *t*, depends only on the environmental variable, *x*, at the lagged monthly interval, *m* ≤ *t*. The FLM was fitted as:

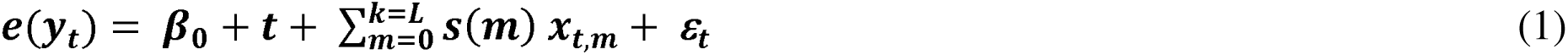

where: ***e*(*y_t_*)** is the response variable estimate, and ***β_0_*** is the model intercept. We fitted the FLMs using a maximum time lag of *L* = 19 months. This 19-month lag length was chosen following an initial model selection procedure comparing different time lags starting from 7 months to 43 months (Appendix S2, Figure S1). The 19-month lag length goes back to December, in the winter, a year and a half before the July growing season census of interest. *C. undulatum* has no seed bank, surviving seeds germinate within their first spring. So, for seedling recruitment, we restricted the maximum time-lag (L) to 11 months, which represents the months since seed release in August of the previous year to germination in June/July of the current year. Since p-values for the smoothing term in generalized additive models are approximate and likely low (Wood, 2017), we only present the results where the overall F-tests for the smooth spline had *p* < 0.01 (Tenhumberg et al., 2018).

When SPEI, precipitation, or temperature were found to have a significant effect, we present them here (Figure 4) (All other effects in Appendix S2, Tables S3, S4). We considered a past weather variable to have a significant effect on a ramet’s vital rate on a census date (month [m] = 0) when the 95% confidence interval did not include the zero line.

We could not estimate several transition probabilities because the FLMs did not converge. These included: four transitions at Arapaho, all involving the inactive stage (IA → SF, IA → IA, IA → MR, and IA → SR; see Figure 1 for stages); and, five transitions at Niobrara (SR → MF, MR → IA, MR → SR, survival of seedlings, and MF recruits), all of which lacked inter-annual variation in their vital rate estimates (Appendix S2, Tables S1, S2).

### 2.7 Predicting ramet population viability under different drought frequency scenarios

In order to examine the viability of *C. undulatum* ramet populations in relation to drought frequency, we used the annual MPMs to perform a population viability analysis (PVA). To do so for each site, we first calculated the annual average SPEI from June of the previous year (*t – 1*) to July of the census year (*t*) using the one-month SPEI values estimated above. Then we sorted the annual MPMs into two groups for each site: years with positive SPEI (non-drought: wetter than normal conditions), and years with negative SPEI (drought: drier than normal conditions).

We started our simulations for each site with a population of 50 ramets, and distributed the 50 ramets according to the stable stage distribution estimated from the mean MPM for the entire study period (1990 - 2009) (Appendix S2, Figure S5). Then, we calculated the ramet population one year at a time into the future by randomly drawing MPMs from each site until we reached 50 years (“project” function in the “popdemo” package, R statistical software).

We ran three scenarios, where frequency of drought: increased, did not change, or decreased. For the increased drought frequency scenario, we sampled 90% of the time from MPM matrices with negative SPEI values and 10% from matrices with positive SPEI. For the no change in drought frequency scenario, we sampled randomly from all MPM matrices. Finally, for the decreased drought frequency scenario, we sampled 10% of the time from MPM matrices with negative SPEI values and 90% from matrices with positive SPEI.

For each of the three scenarios at each site, we repeated the simulation 1,000 times. Each simulation resulted in a different ramet population size after 50 simulated years. We plotted the distribution log median ramet population sizes and calculated how likely the simulated population size after 50 years was lower than the starting population size. A decreasing population size would indicate a nonviable population.

## 3.0 Results

### 3.1 Demography: Numerical Patterns and Mean Vital Rates by Site

#### Ramet Numbers

Over the 20 years, 9,053 ramets were observed at Arapaho and 3,299 at Niobrara. At Arapaho, the average annual ramet density per plot of 113.2 (SE: 9.72) was significantly higher than that at Niobrara, 55.0 (SE 2.63) (Table 1; t = −5.16, *p* < 0.01). At both sites, the predominant ramet stage was juvenile single rosette (Table 1): 81.3% of all ramets at Arapaho and 83.5% at Niobrara. The next most common ramet stage was: flowering single rosettes at Arapaho (7.4%), and seedlings at Niobrara (6.1%). All other stages were less than 5% (Figure 1).

**TABLE 1.**
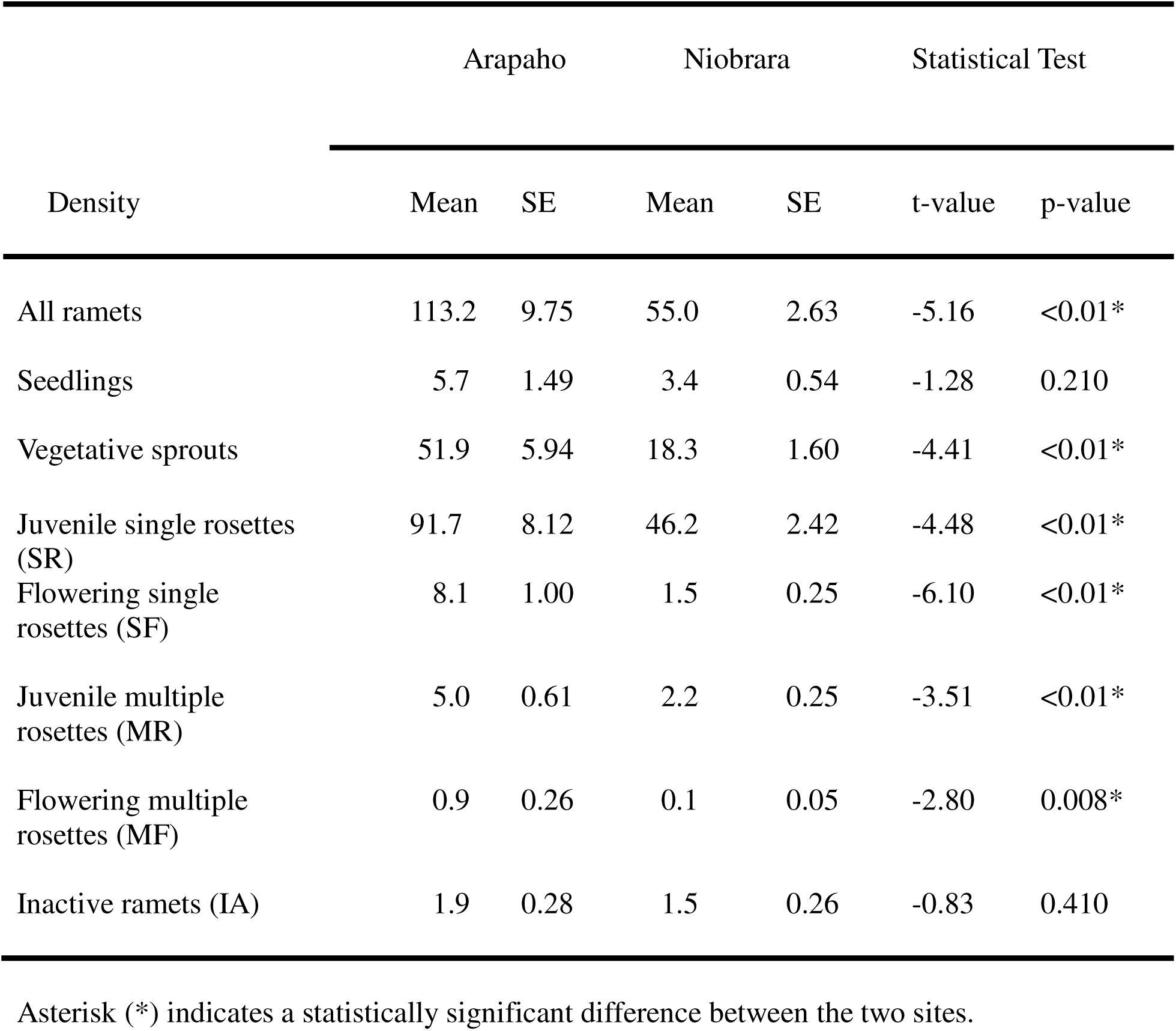
Mean annual ramet density of *C. undulatum* per 12 m x 12 m plot (1990 - 2009) by site (Arapaho: N = 4 plots/year; Niobrara: N = 3 plots/year).

#### Survival

We observed a few ramets (<0.01%) live 11 years without flowering. However, most ramets died within their first two years and did not flower (Arapaho 73.2%; Niobrara 66.2%). At both sites, ramets with multiple sub-rosettes had the highest annual probability of surviving (Arapaho 72.9%; Niobrara 77.4%), higher than those in the dominant single rosette stage (Arapaho 65.3%; Niobrara 64.9%) (Table 2). The difference in survival between multiple and single rosette ramets was significant at both sites: Arapaho (t = 2.95, *p* = 0.008) and Niobrara (t = 2.35, *p* = 0.03). Seedlings had the lowest annual survival rate among stages at both sites (Arapaho 50.6%; Niobrara 47.3%) (Table 2).

**TABLE 2.** Mean transient and asymptotic growth rate (λ) and survival for the vegetatively active ramets (seedling, single rosette, multiple rosette).

|  | Arapaho |  | Niobrara |  | Test Statistics |  |
| --- | --- | --- | --- | --- | --- | --- |
|  | Mean | SE | Mean | SE | t-value | p-value |
| Asymptotic ($\lambda_t$) | 1.22 | 0.086 | 1.05 | 0.045 | 1.25 | 0.22 |
| Transient | 1.25 | 0.17 | 1.03 | 0.043 | 0.27 | 0.78 |
| *Survival of:* |  |  |  |  |  |  |
| Seedling | 0.51 | 0.06 | 0.51 | 0.06 | 1.07 | 0.28 |
| Single rosette | 0.66 | 0.02 | 0.66 | 0.03 | 1.37 | 0.19 |
| Multiple rosette | 0.74 | 0.04 | 0.78 | 0.85 | 0.85 | 0.39 |

#### Flowering

The mean annual proportion of ramets that flowered was quite low (Arapaho 8.1%; Niobrara 3.1%). Most flowering ramets were single rosetted (Arapaho 90.2%; Niobrara: 92.0%) rather than multiple rosetted (Arapaho 9.8%; Niobrara 8.0%). Most flowering ramets transitioned from single rosettes the previous season (Arapaho 74.4%; Niobrara 76.8%); however, a small portion of flowering ramets were recruited as sprouts from unknown taproots (Arapaho 17.1%; Niobrara 9.5%).

#### Recruitment

Each year, on average, new ramets (seedlings and sprouts) composed about half of those observed (Arapaho 49.4%; Niobrara 57.1%) (Table 1). The average density of seedling recruits per plot was low: 5.7 (SE 1.49) at Arapaho and 3.4 (SE 0.54) at Niobrara (Figure 2B). Despite the variation among years, these means did not differ significantly (t = −1.28, *p* = 0.21). At both sites, most new ramets recruited each year were vegetative sprouts: 90.1% at Arapaho, 84.3% at Niobrara. Ramet sprout recruits averaged 51.9 (SE: 5.94) per plot at Arapaho (Figure 2C), which was significantly greater than the surprisingly low average of 18.3 (SE 1.60) per plot observed at Niobrara (t = −4.41, *p* < 0.01).

**FIGURE 2.**
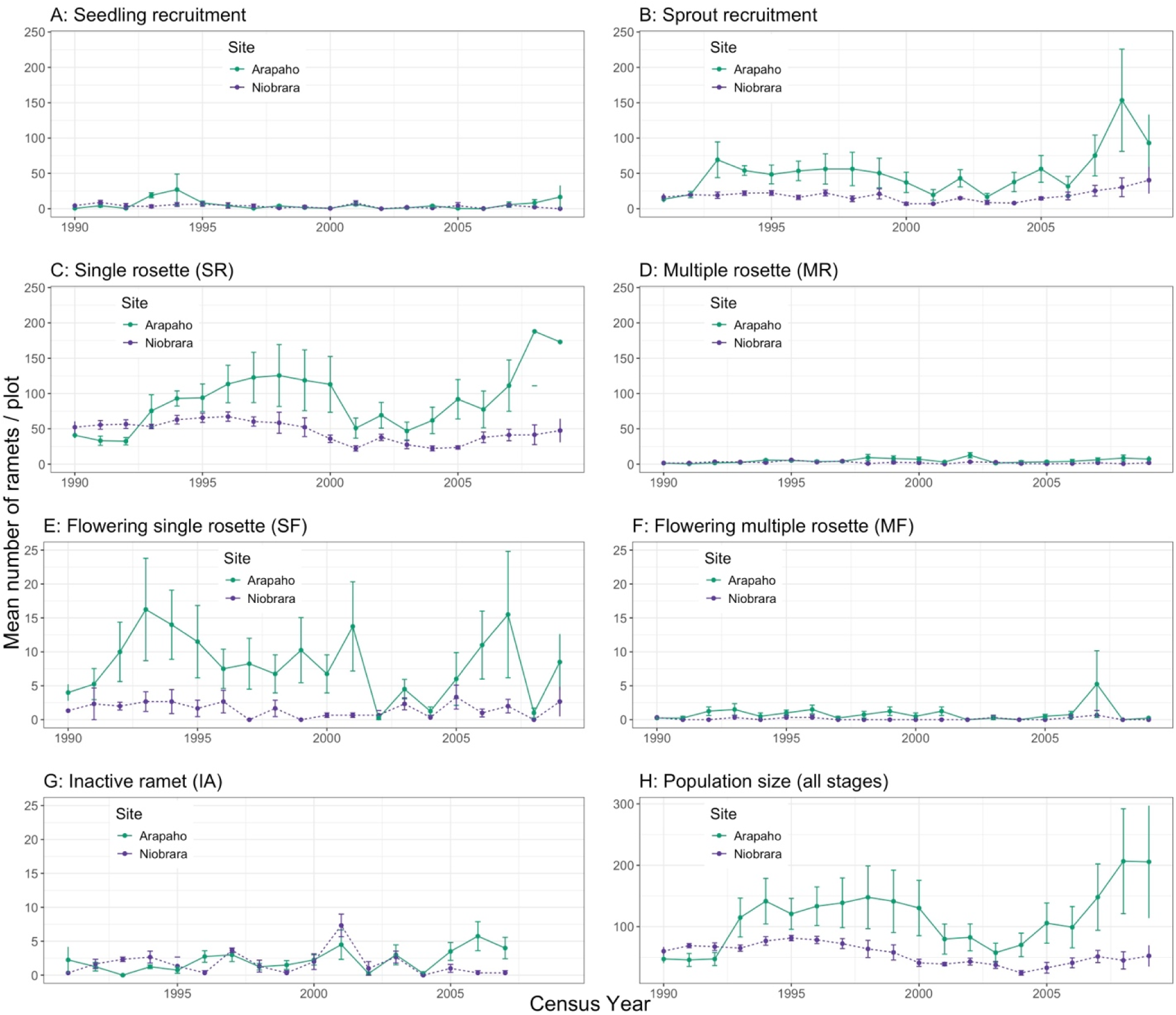
Mean (SE) number ramets per plot by stage for each site: (A) seedling recruits, (B) total vegetative sprout recruits, (C) juvenile single rosettes, (D) juvenile multiple rosettes, (E) flowering single rosettes, (F) flowering multiple rosettes, (G) inactive ramets, and (H) total ramet population size. Management differed between sites. At Arapaho, plots were hayed in late July after the population census every four years, while at Niobrara, plots were lightly grazed, primarily in early season (Details in Methods).

### 3.2 Matrix Population Model

At both sites, the asymptotic and transient growth rates were similar. At Arapaho, the average annual asymptotic growth rate (λ_t_) was 1.22 (SE: 0.09), and the observed average transient growth rate was 1.25 (SE: 0.17) (t = 1.07, *p* = 0.29; Figure 3A). At Niobrara, the average annual asymptotic growth rate (λ_t_) was 1.06 (SE: 0.045), and the observed average transient growth rate was 1.03 (SE: 0.04) (t = 0.955, *p* = 0.35; Figure 3B). Although the average asymptotic growth rate at Arapaho (1.25) appeared to be higher than that at Niobrara (1.06), the difference was not significant (t = −1.77, *p* = 0.85).

**FIGURE 3.**
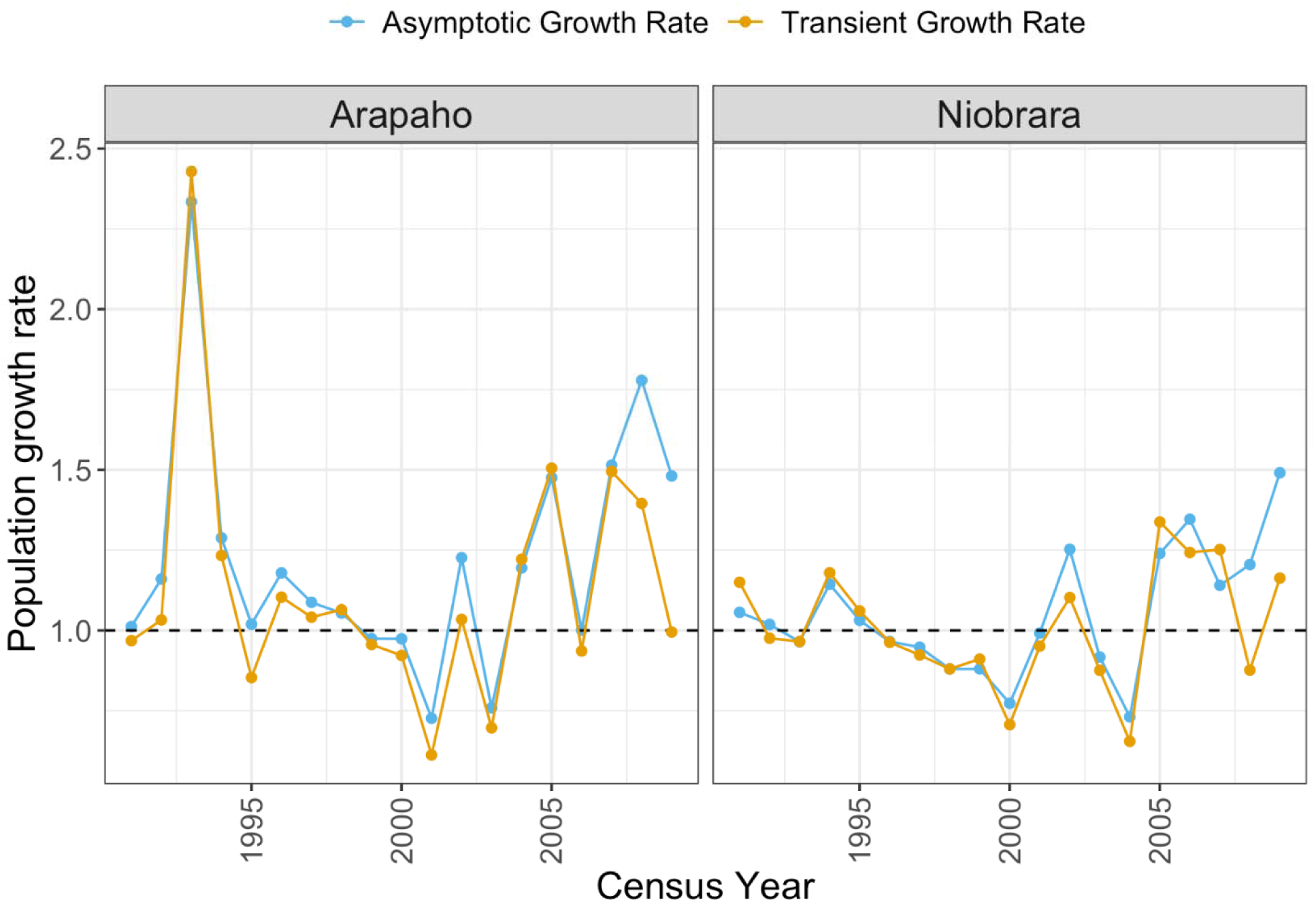
Variation in estimates of asymptotic (blue) and transient (orange) population growth rates over the study by site; asymptotic growth rate was estimated from the Matrix Population Model (MPM) constructed from the field data.

The annual population growth rate varied at both sites during the study, with most of the 19-year transitions showing an increase in population (Arapaho = 68.4%; Niobrara = 57.9%). In 1993, at Arapaho, we found a spike in the population growth rate (Figure 3A), reflecting exceptionally high recruitment in 1993; this led to a 59% increase in population size in 1993 over 1992.

### 3.3 Temporal variation in weather conditions

We defined the baseline period for weather variables as the 40 years (1950 - 1989) prior to the beginning of the demographic census in 1990. Then, in order to allow examination of up to a 19-month lag in weather parameters for examining demographic responses, we included 1988 and 1989 with the 1990 – 2009 study period.

#### Difference in weather conditions between sites

The mean annual temperature observed was similar at the two sites from 1988 - 2009 (Arapaho = 8.93°C, SE = 0.60, Niobrara = 8.98°C, SE = 0.80, t = 0.05, *p* = 0.96; Figure 4A). Mean annual precipitation, however, was significantly lower at Arapaho (39.1 mm, SE = 2.19) than at Niobrara (47.2 mm, SE = 2.62) (t = 3.20, *p* = 0.002). The one-month SPEI values, evaluating monthly drought conditions, were significantly higher (wetter) at Niobrara than at Arapaho (t = 13.36, *p* < 0.01).

**FIGURE 4.**
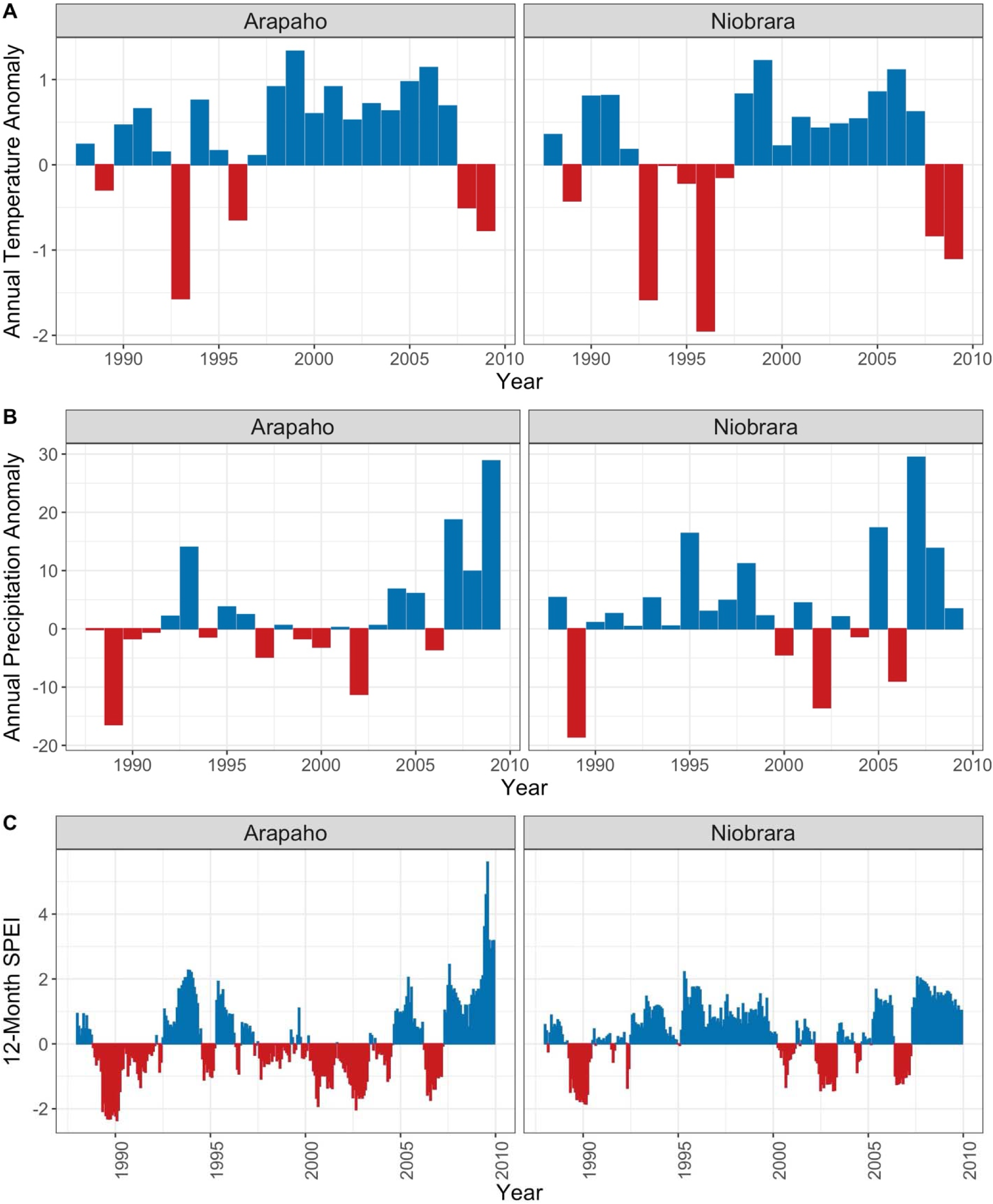
Variation in weather variables during the study in: (A) annual temperature anomaly (°C), (B) annual precipitation anomaly (mm), and (C) 12-month Standardized Precipitation Evapotranspiration Index (SPEI) by site. Anomaly is defined as annual deviation from baseline reference conditions (1950 – 1989). Blue bars depict warmer or wetter than baseline conditions; red bars depict colder or drier than baseline conditions.

#### Change in weather conditions compared to baseline

Average annual temperatures during the study period were somewhat higher than during the baseline period at both sites, by 3.8% at Arapaho and by 1.4% at Niobrara (mean 8.6°C). Average annual precipitation during the study also was higher during the study than during the baseline period at both sites: 6.1% above the 36.9 mm baseline average at Arapaho, and 7.9% above the 43.7 mm baseline average at Niobrara.

### 3.4 Time-lag effect of weather variables on vital rates and population growth rates

#### Overall Model Outcome

To explore the effect of variation in past weather conditions on subsequent population parameters, we examined lagged effects of current deviations from the reference period, in monthly mean temperature anomaly, total monthly precipitation anomaly, as well as in one-month SPEI.

First, lags in monthly mean temperature significantly affected only one vital rate at one site, Arapaho: the transition from single rosette (SR) to flowering multiple rosette (MF) (Figure 5E). None of the rates at Niobrara were affected by temperature at any lag interval (Details: Tables S3&S4).

**FIGURE 5.**
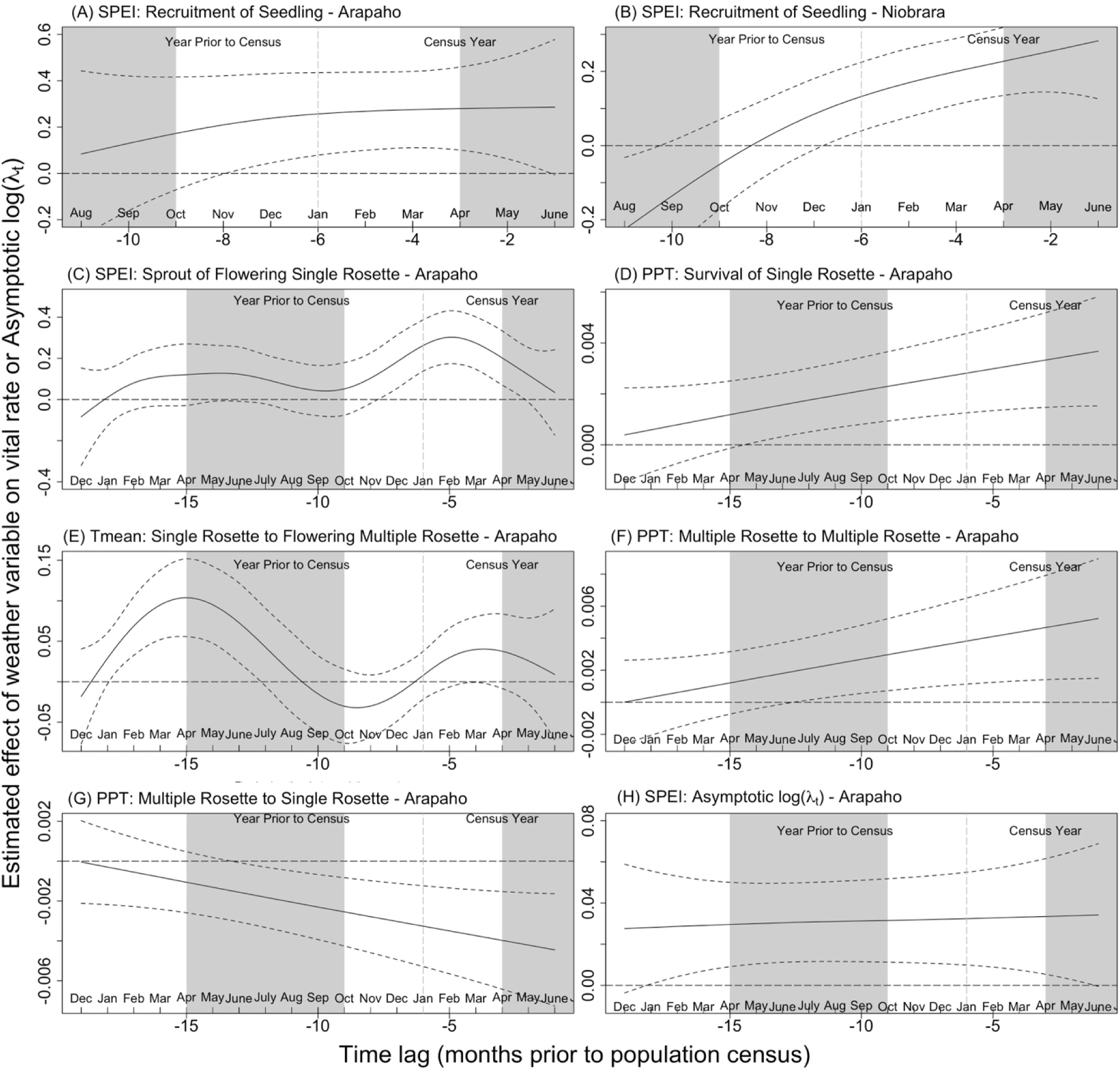
Significant time-lagged effects of the three weather variables on vital rates and on asymptotic growth rate (λ_t_) over the 20 years of the study (non-significant effects: Table S3&S4): effect of SPEI on: seedling recruitment at Arapaho (A) and at Niobrara (B), vegetative sprouts recruited as flowering single rosettes at Arapaho (C), and log asymptotic population growth rate (log λ_t_) at Arapaho (H); and, effect of precipitation at Arapaho on: survival of single rosettes (D), stasis of multiple rosette stage (F), retrogression (shrinkage) of multiple to single rosettes (G); and, finally, effect of temperature on transition of single rosettes to flowering multiple rosettes at Arapaho (E). Vertical gray areas indicate growing season (May - September), and white areas indicate the non-growing season (October - April). The X-axis represents the discrete monthly time lag prior to the July census, from month (m = 0) in July on the right back to December two years prior (m = −19) on the left; however, for seedling recruitment from prior year seed production the maximum lag was 11 months, given the lack of a seedbank. The lines indicate: horizontal dashed line at zero = no weather effect; solid line = mean effect of the weather variable with time; and, dotted lines = 95% confidence interval around the mean. We defined time lag to be significant when the 95% confidence interval did not overlap with the horizontal dashed "no effect" line.

Second, time-lags in total monthly precipitation had a significant effect on four vital rates, all at Arapaho: seedling recruitment, single rosette survival, multiple rosette to multiple rosette stasis, and multiple rosette reversion to single rosette transition (Figure 5D, 5F, 5G). At Niobrara, the wetter site, no significant lagged effects of variation in precipitation were observed on any stage transitions or other vital rates (Details: Table S3).

Finally, lags in SPEI (a measure of drought) affected four demographic parameters, three at Arapaho and one at Niobrara. At Arapaho, lags in SPEI effects were significant for two ramet vital rates, recruitment of seedlings and recruitment of single flowering rosettes via sprouting, and for the asymptotic population growth rate (log (λ_t_)) (Figure 5A, 5C, 5H) (Table S3). At Niobrara, lags in SPEI effects were significant for only one vital rate: seedling recruitment (Figure 5B; Table S4).

#### Recruitment

Seedling recruitment was greater when SPEI was wetter-than-average (positive SPEI) in the interval between seed release and subsequent establishment. At Arapaho, seedling recruitment was highest when SPEI was wetter-than-average starting 8 months prior to the growing season, from November of the previous year until the census date, *t* (Figure 5A). At Niobrara, seedling recruitment was higher when the weather was wetter than average (positive SPEI) starting 7 months prior, from December of the year before the census (*t* – 1) until the census date, *t* (Figure 5B). So, a wet winter and spring before the census season increased seedling recruitment at both sites.

At Arapaho, but not at Niobrara, the number of vegetative sprouts recruited as flowering single rosettes (SF_rec_) was higher if it was relatively wet (positive SPEI) from eight to two months before the census, from November of the previous year (*t* – 1) to May of the census year (*t*) (Figure 5C). We did not find a significant effect of SPEI on the numbers of vegetative sprouts recruited as other stage at either site (Table S3&S4).

#### Survival

At Arapaho, but not at Niobrara, survival of single rosettes, the numerically dominant stage, was higher when higher-than-normal precipitation occurred, starting 14 months prior to the census (May in *t* – 1) until the July census date (July in *t*) (Figure 5D). However, the absolute magnitude of the precipitation effect was small since the slope of the precipitation line was close to zero (CI: 0 - 0.004; Figure 5D). We did not find a significant lagged effect of weather variables on other variables reflecting survival, such as survival of seedlings or of multiple rosettes, at either site (Table S3&S4).

#### Flowering probability

We found two effects of the weather parameters on flowering probability at Arapaho, but none at Niobrara. First, vegetative sprouts emerging as single flowering ramets were increased with wetter-than average (positive SPEI) from 7 months to 2 months before the July census (Figure 5C). Second, single rosettes are more likely to transition flowering multiple rosettes when temperatures were warmer-than-average from 18 months to 12 months prior to census, from January to July of the year before the census (t - 1) (Figure 5E). We did not find a significant effect of any of the weather variables on any of the other flowering transitions at either site (Table S4).

#### Stage transitions

Most stage transition rates at both sites were unaffected by the variation in the weather variables measured (Table S3&S4). However, at Arapaho, but not Niobrara, the fate of the multiple rosette stage was influenced by precipitation. More ramets stayed in the multiple rosette stage (stasis), rather than transitioning to another stage, when higher-than-normal precipitation occurred in the year before census, specifically starting 12 months before census (July of *t*-1) until the census date (July of *t*) (Figure 5F). Further, more multiple rosette ramets shrank, transitioning to single rosettes, when lower-than-normal precipitation occurred in the year before the census, starting 13 months (June in *t* – 1) to the census date (July in *t*) (Figure 5G). The magnitude of the effect of these variations in precipitation, however, was small for both of these transitions since both slopes were close to zero (CI: 0 - 0.006; Figure 5F, 5G).

#### Population growth rate, log (λ_t_)

At Arapaho, but not at Niobrara, we found that the asymptotic population growth rate, log (λ*_t_*), was highest when the census was preceded by a relative wet period (positive SPEI), starting 18 months before census (from January of *t* – 1 to June of census year *t* (Figure 5H). Again, however, the absolute effect size was small since the SPEI slope was close to zero (CI: 0.0 - 0.06).

At Arapaho, the estimated mean and variation of annual asymptotic population growth rates (λ_t_) were significantly higher during wet years (positive SPEI) (mean λ_t_ = 1.47, range of λ_t_ = 1.09 - 2.33) than dry years (negative SPEI) (mean λ_t_ = 1.03, range of λ_t_ = 0.73 - 1.48) (t = 3.08, *p* = 0.006; Figure 6A). Conversely, at Niobrara, the estimated mean and variation of λ_t_ were similar during wet years (positive SPEI) (mean λ_t_ = 1.05, range of λ_t_ = 0.77 - 1.49) and dry years (negative SPEI) (mean λ_t_= 1.05, range of λ_t_= 0.73 - 1.35) (t = 0.031, *p* = 0.97; Figure 6B).

**FIGURE 6.**
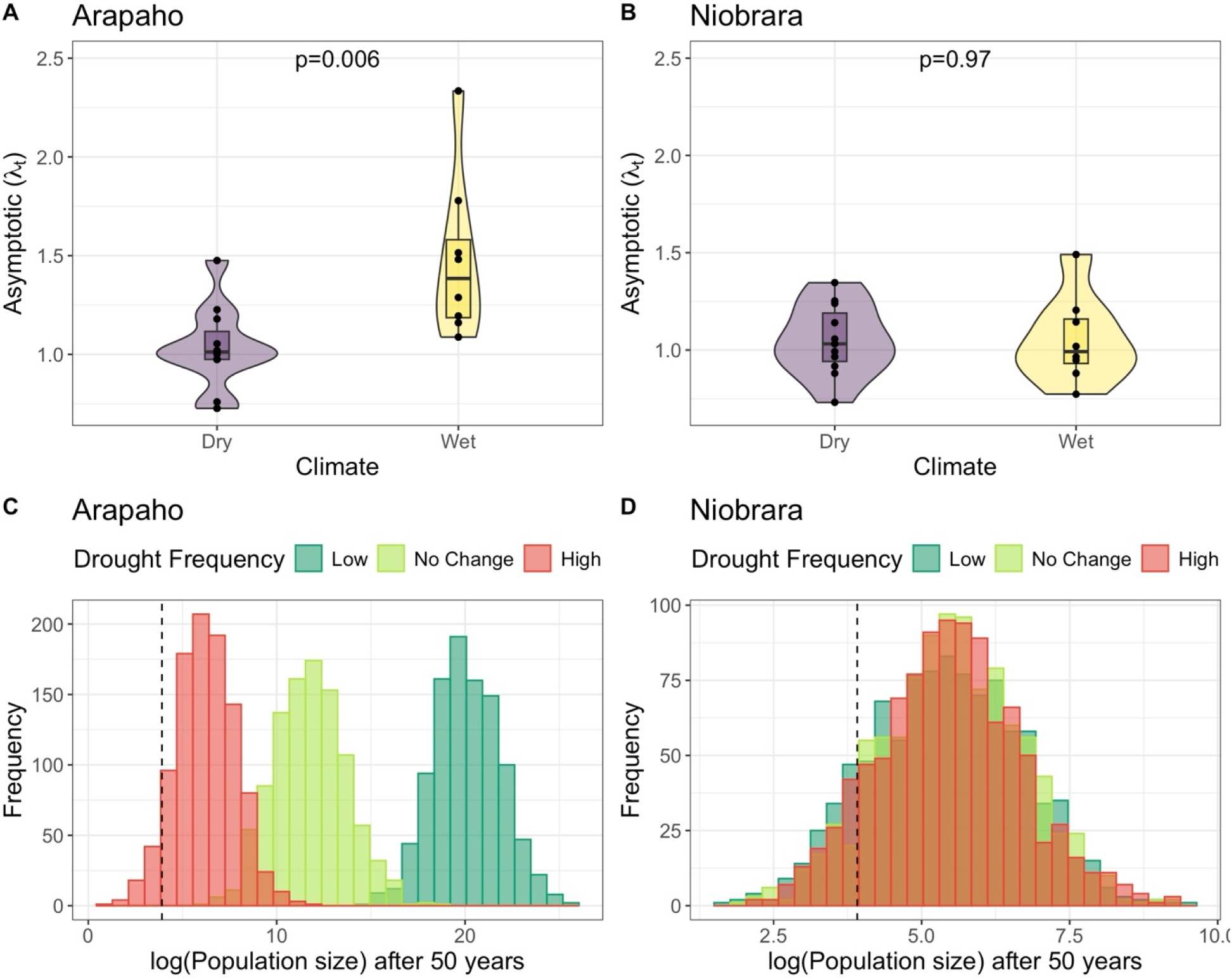
(A, B) Distribution of asymptotic population growth rate (λ_t_) from the projection model by site during wet and dry years; the outlines of the violin plot represent the kernel probability density, meaning the width of the shaded area indicates the proportion of data in that range. (C, D) Distribution of simulated population size after 50 years at each site under high, normal, and low drought. We defined drought frequency as follows: “high” meant drier, with the population experiencing dry years 90% of the time; “low” meant wetter, with the population experiencing dry years only 10% of the time; and “normal/no change where we randomly picked matrices from the pooled matrices for each site (Details in Methods Vertical dashed lines are the starting population (50 ramets), and the bars are the frequency distribution from 1000 simulations.

### 3.5 Ramet population size under three drought scenarios

We simulated the change in population size of *C. undulatum* at the two sites during different drought scenarios over 50 years. First, at Arapaho, under the assumption drought frequency does not change, the simulated population size increased from an initial 50 to a median of 125708 ramets; not a single run out of 1,000 predicted a decrease in population size (dark green bars in Figure 6C). Alternately, under the assumption drought frequency increases (90% MPMs associated with dry years), the median simulated population size was only 488 ramets; in 6.8% of the 1,000 runs, the projected population size was below the starting population (red bars in Figure 6C). Under the assumption that drought frequency decreases (10% MPMs associated with dry years), the median simulated population size was 487,842,000 ramets (light green bars in Figure 6C).

At Niobrara, the distribution of simulated population sizes overlapped for all three drought scenarios; median population size was 219, 249, and 232 ramets for low, unchanged, and high drought frequency (Figure 6D). This result was logical, given that the average λ_t_ values were similar for wet and for dry years (Figure 6B). Out of 1,000 runs, 13.1%, 8.5%, and 10.4% of the runs were projected to decrease below the starting population for low, no change, and high drought scenarios, respectively (Figure 6D). Overall, the predicted median population size under no change in drought scenarios was significantly lower at Niobrara than at Arapaho (t =-214, *p* < 0.01).

## 4.0 Discussion

### 4.1 Site differences in *C. undulatum* ramet demography

In this study, we analyzed the dynamics of *C. undulatum* ramet populations using a 20-year-long demographic data set collected at two regions within the sandhills prairie grasslands in Nebraska. The weather data showed that Arapaho is relatively drier than Niobrara, and because water availability is typically the primary limiting factor for plant productivity in the Nebraska Sandhills (Chaves et al., 2003; Louda, 2000; Schwinning & Sala, 2004), ramet populations might be expected to be higher at Niobrara. However, we found the opposite; *C. undulatum* ramet density was higher at Arapaho than at Niobrara.

The differences in ramet density between the sites 270 km apart in the Sandhills could reflect site-specific demographic responses to growing conditions by *C. undulatum.* The population at Arapaho had higher vegetative sprout recruitment rates and a higher proportion of ramets flowering than did the population at Niobrara. Also, even though more ramets flowered at Arapaho, seedling establishment was similar between sites; so, suggesting either the drier environment at Arapaho reduced the germination rate (e.g., Markesteijn & Poorter, 2009) or pre-germination seed loss was higher there (Rand et al., 2020). Significant pre-dispersal seed predation on native thistle flower heads has been recorded at both sites (Louda & Potvin, 1995; Russell et al., 2010; West & Louda, 2021), and the added unexpected losses imposed by the biological control weevil, *Rhinocyllus conicus,* have led to a decline in the population density of a co-occurring native congener, *Cirsium canescens* (Louda, 2000; Rand et al., 2020). However, such pre-dispersal seed predation is likely less important in the *C. undulatum* population density because, unlike *Cirsium canescens*, vegetative recruitment by *C. undulatum* occurs via asexual shoots from the perennial taproot (Figure 2B; Kaul et al., 2011).

In addition, some of the difference in densities likely reflects the fact that the study areas were managed differently. At Niobrara, early and late season cattle grazing annually likely reduced flowering success (De Bruijn & Bork, 2006). At Arapaho, mowing occurred only at four year intervals, in late summer after flowering and seed set. Both management strategies are standard practices to maintain grassland productivity (Tälle et al., 2016). However, depending on the timing of the management action, they can have an effect on plant populations (Lennartsson et al., 2012; Nakahama et al., 2016). Since our study was not designed to evaluate the effect of different prairie management on the dynamics of *C. undulatum*, it is unclear what contribution, if any, management had to the observed differences in flowering and density between sites.

### 4.2 Influence of weather variables on *C. undulatum* ramet demography

We assessed the effect of temperature, precipitation, and SPEI on ramet vital rates and asymptotic population growth rate (λ_t_) of *C. undulatum* ramets. At Arapaho, we found significant correlations between temperature, or precipitation, or SPEI and six out of 18 vital rates and on λ_t_; however, at Niobrara, only one out of 17 vital rates was affected significantly. Three hypotheses emerge to explain the lower number of vital rates affected significantly by weather variables at Niobrara compared to Arapaho. First, Niobrara was wetter overall than Arapaho (Figure 3), generally moderating drought and variable conditions. Second, Niobrara had smaller ramet population sizes (Figure 2) (Austin & Leckie, 2018), and these could have reduced our statistical power to detect weather effects (Lindell et al., 2022). Lastly, the 20-year time series might still be too short to have sufficient power to detect any subtle effects of weather on demographic rates at Niobrara. For instance, (Teller et al., 2016) suggested that one may need at least 25 – 30 years of observational data to detect a significant delayed effects of weather variables, although we did detect such effects at Arapaho in the 20-year time frame. In the following paragraphs, we discuss the lagged effects of weather variables detected on key vital rates.

#### Recruitment

Our study suggests that in both sites, seedling recruitment was highest when the winter and spring months preceding the population census were wetter than normal (positive SPEI), likely increasing germination success. Wetter than normal conditions provide *C. undulatum* seeds with the required moisture level for germination. Adequate soil moisture is critical for seed imbibition and metabolic activation as a critical first step in thistle seedling establishment (Eckberg et al., 2017).

In addition, vegetative sprout recruitment varied. At Arapaho, the drier of the two sites, but not at Niobrara, flowering single rosettes that appeared for the first time via vegetative sprouts increased when the winter and spring months preceding the month of May of the census season were wetter than normal. The observed delayed effect of prior wet conditions on rapidly flowering sprouts may be associated with bud bank development. An increased moisture in the form of snow during the preceding winter season may increase the number of viable buds available to sprout, or replenish soil moisture enough to stimulate flowering initiation, as well as insulate the bud bank from frost damage (Evers et al., 2021; Schwinning & Sala, 2004). Further, wetter conditions during the spring, early in the growing season, may facilitate the growth of the underground plant structures (Dalgleish et al., 2011).

#### Survival

At drier Arapaho but not Niobrara, we found that the survival of single rosettes, the predominant stage, was increased by the cumulative effect of precipitation from the previous year’s summer until the current growing season. Precipitation during the growing season facilitates photosynthetic activity and, hence, resource production and accumulation, promoting growth (Song et al., 2016); and, larger plants are more likely to survive in the next growing season (Atkinson et al., 2022).

#### Flowering probability

In addition to the vegetative recruitment of flowering ramets (see above), at Arapaho, but not Niobrara, we also found that the probability of transitioning from a single rosette to a flowering multiple rosette increased if the previous year’s winter, spring, and summer temperatures were high. Other studies found a similar delayed effect of temperature on flowering probability; delayed flowering correlated with higher summer temperature from one year ( *Pulsatilla vulgaris*; Lindell et al., 2022), from two years (*Veratrum tenuipetalum*; Iler & Inouye, 2013), or from four years (*Frasera speciosa*; Evers et al., 2021) prior to inflorescence emergence. If the previous year’s growing season is warmer, photosynthetic and photorespiratory rates may increase rapidly (Gu et al., 2022). Such a response may result in larger resource accumulation in the root system of *C. undulatum* available for developing multiple rosettes and bolting stems.

#### Stage transition

At Arapaho but not Niobrara, the transition from multiple rosette to multiple rosette stage (stasis) increased when the precipitation was high from the summer of the previous year to the population census. Field observations suggest that the development of a multiple rosette by *C. undulatum* is triggered by damage to the apical meristem, often by insect feeding (Svata Louda *pers com*); this releases the shoot from apical dominance. We hypothesize that high precipitation, and hence higher resource accumulation, facilitates ramet growth and survival of sub-rosettes of a multiple rosette.

Comparably, at Arapaho but not Niobrara, when precipitation was low from the previous summer to the census data, multiple rosette ramets (MR) regressed to the single rosette (SR) stage. Under a prolonged period of low precipitation, a multiple rosette may not have enough resources to sustain multiple sub-rosettes; if so, it may allocate the available limited resources to the largest sub-rosette and let the others die (Ehrlén & Van Groenendael, 2001). This hypothesis is consistent with the suggestion that plants endure extreme conditions, such as drought, by reducing their overall biomass, or “shrinking,” to conserve resources (Vega & Montaña, 2004).

#### Population growth rate, log(λ_t_)

At Arapaho but not Niobrara, we found that log(λ_t_) was high if *C. undulatum* ramets experienced wet conditions (positive SPEI) during the previous 18 months. This result aligns with many other studies that show water availability is critical in semiarid systems and, thus, plant populations should perform better under increased water availability (Dalgleish et al., 2011; Evers et al., 2021; Stephenson et al., 2019).

### 4.4 Predicting *C. undulatum* ramet population viability under three different drought frequency scenarios

Matrix Population Models (MPMs) are powerful tools for assessing population viability (Crone et al., 2011). In this study, we first examined the differences in estimated λ_t_ values between wet years and dry years. We found that at Arapaho, but not Niobrara, λ_t_ values were significantly greater in wet years than in dry years (Figure 6A, B). Thus, using the parameters observed and changing drought frequency influenced simulation outcomes at Arapaho (Figure 6C), but not at Niobrara (Figure 6D).

At Arapaho, the average annual population growth rate (λ_t_) was > 1 during both wet and dry years at both sites, suggesting the population is viable and potentially growing. However, in the presence of environmental stochasticity, it is possible that a run of “bad luck” leads to a population size after 50 years that is smaller than the initial population size, especially if the drought frequency increases. By randomly selecting transition matrices from our observed MPMs for the 20 years, we estimated the effect of environmental stochasticity on the change in population size (Menges & Dolan, 1998).

Our simulations suggest that at Arapaho it is highly likely that *C. undulatum* is viable for the next 50 years, under all three drought scenarios - current, decreased or increased drought frequency. Interestingly, even when drought frequency increased, the simulated ramet population expanded after 50 years in 93% of runs. At Niobrara, the simulated outcomes were similar under all three drought scenarios, consistent with the dry-vs-wet comparison, with population increased in 87% of the 1,000 runs. Thus, we conclude that *C. undulatum* ramet populations at both sites are likely viable for the next 50 years at Niobrara (Figure 6C, 6D), even in the context of predicted climate changes.

## Acknowledgments

This study was done on the homelands of the Lakota People (Arapaho Prairie Preserve) and of the Ponca People (Niobrara Valley Preserve); we honor this legacy. We are also indebted to The Nature Conservancy, Nebraska Chapter, for permission to work in these reserves and to their incredible staff for all of the support and encouragement provided for the study, especially A. A. Steuter at the beginning of the study. The field data were collected by S. M. Louda with the help of her collaborators: A. Arnett, T. A. Rand, F. L. Russell, R. W. Otley, 18 graduate students and 21 undergraduate students over the 20 years. Our thanks and gratitude to all these wonderful lab and field crew members – we could not have done the work without you; your conscientious, cheerful, committed involvement and assistance made a difference. The field work was supported in part by several grants for thistle research to S.M.L.: N.S.F., Division of Environmental Biology, Ecological Studies, Ecology Program grants DEB96-15299 and DEB04-14777; U.S.D.A. National Research Initiative, Biology of Weedy and Invasive Plants OEP2000-0088; The Nature Conservancy, C. A. Ordway and L. Robinson Research Grant 1997-2000; U.N.L., School of Biological Sciences Research Grants; and, these grants were augmented by personal funds.

## Authors Contribution

SML is responsible for the design and collection of the field data and initial data management. KHK is responsible for the extraction of ramet survival numbers along with subsequent data management, including error checking. JNM, with guidance and assistance from BT, developed the models and did the statistical analyses. JNM wrote the original manuscript draft. All authors reviewed and approved the final version of the manuscript.

## Data availability statement

The demographic data associated with this study can be accessed here: https://doi.org/10.6084/m9.figshare.29019173.v1. The R code associated with data analysis is available here: https://github.com/JohnMensah50/Ramet_Delay_Analysis.

