## Appendix S1 for "Delayed effects of water-limiting conditions influence the ramet demography of a native iterocarpic thistle"

**1.0 Data management for estimating annual transition rates**

This section provides an overview of the data management and cleaning process involved in processing the original data from the field to an analysis-ready format appropriate for our study. The R code associated with the data cleaning is available on GitHub (https://github.com/JohnMensah50/Ramet_Delay_Analysis).

In 1990 – 1999, ramets in the plots were monitored twice a year: in early season (May) and in late season (late June - mid July); plots at both sites were examined within one week each year. For these years, we used the late-season observations when available, since they represented full season performance, including emergence and flowering; incomplete emergence was observed in May and flowering effort was incomplete, compared to June-July observations. Our rules were as follows, and the few exceptions to use of July data are explained below:

A. If the stage in the late observation was the same as the stage in the early observation, then that stage was unambiguous and the data associated with the late observation was used for that year.
B. If the stage in the late observation was different from the stage in the early observation, then we used the late observation as the stage for the year, with three relatively infrequent exceptions:
    1. If the ramet had no record (was missing) in the late-season observation, we used early-season record; we assumed that early data was better than no information since it maintained evidence on all vegetatively active ramets.

2. If the ramet was recorded as a seedling in the early season, then we recorded as a seedling for that year, whether the cotyledon were present or not by the late season observation.
3. If the stage for the late observation stage was blank (missing), we used the stage recorded in early season, as the best information available, and it also helped maintain information on vegetatively active ramets.

In 4 years (2000 - 2004), all ramets were measured in late May, but in general only flowering ramets were remeasured in July; however, some late-season observations were recorded on non-flowering ramets. To standardize, we removed all late-season observations that were not flowering ramets. Then we joined the late-season data on flowering ramets to their early season data along with the non-flowering ramets measured in early season. Finally in 2002, data on all ramets were collected in May and in June (mid-season); we treated this year using the same rule used to merge early and late-season observations 1990-1999 above.

In 5 years late in the study (2005 – 2009) the ramets were measured only once, in mid- to late-season: in mid-July for 3 years (2005 - 2007) and in late June for two years (2008 - 2009). Since these were mid-/late-season, we used these data as collected. Ramet sprouts in 2008 – 2009 at the end of the study were no longer given numerical tags; so, those did not contribute to estimation of specific stage transition rates.

### 2.0 CALCULATION OF DEMOGRAPHIC TRANSITIONS

We estimated annual transition rates between the ramet life history stages in late season, specifically July in year t to July in year t+1. To estimate these rates for each ramet stage, including survival and recruitment stages, we used Generalized Linear Mixed Models (GLMMs), with year as the random effect ("glmer" function in lme4 package) (as in Tenhumberg et al.

2018). Each estimated annual vital rate was extracted from the coefficient of random effect estimate, using the *"coef"* function in R statistical software. In the few cases where no transition occurred in a particular year, we substituted the estimated overall mean (the fixed effect of the model) for the missing annual estimate.

***Survival Rate.*** All inactive ramets were treated as surviving, reflecting the definition of an "inactive" ramet as one that reappeared as a sprout at its tag in less than three years. Those that did not reappear were assumed to have died in their year of disappearance.

For the other vegetative ramet stages, *y,* including seedling, single rosette, or multiple rosette, the probability of surviving from year *t* to *t*+1, was conditioned on the ramet not having flowered ($F_t$) in stage *i* in the previous year *t,* since flowering ramets of *C. undulatum* die after flowering. The conditional term of not flowering, which was represented as "$|not\ F_t$" at the left-hand side of equation 1 (below), was used to account for the death of ramets from all causes other than the fatal effect of flowering.

In the model for survival probability (Equation 1), "**logit**" refers to the logit function, which is the logarithm of the odds of an event occurring. And **e** represents the expected value; $\boldsymbol{surv_{i,y,t+1}}$, which denotes the survival probability of individual *i* in stage *y* in year *t+1.* Also, $\boldsymbol{|not\ F_t}$ means conditional on not having experienced flowering event in year t, while $\boldsymbol{\beta_0}$ is the intercept term in the model. Finally, $\boldsymbol{(1|Year)}$ indicates a random effect for the year, allowing for year-specific variations in the intercept. Thus, the model for survival probability is given as:

$$logit[e(surv_{i,y,t+1}|not\ F_t)] = \beta_0 + (1|Year) \quad (1)$$

$$surv_{i,y,t+1} \sim Bernoulli[e(surv_{i,y,t+1})]$$

***Stage transition probability*.** We estimated the transition from ramet stage *x* (*x* ∈{single rosette, multiple rosette, or inactive}) in year *t* to stage y in year *t*+1 (*y* ∈{single rosette, multiple rosette, or inactive ramet}) using Equations 2, 3 and 4, depending on specific stage transition as follows. We conditioned the probability of a ramet *i* transitioning to either a single rosette (SR) or a multiple rosette (MR) (Equation 2) on whether the ramet *i* from stage *x* survived in year *t*+1 and was neither flowering (F) nor inactive (IA) in year *t*. Thus, the model for the annual transition probability of a ramet *i* in *t* to either SR or MR ramet in *t+1* is given as:

$$logit[e(x_{i,t} \rightarrow y_{i,t+1} | surv_{i,x,t+1}, not\ IA_t not\ F_t)] = \beta_0 + (1|Year) \quad (2)$$

$$x_{i,t} \rightarrow y_{i,t+1} \sim Bernoulli[e(x_{i,t} \rightarrow y_{i,t+1})]$$

We conditioned the probability of ramet *i* in stage *y* in year *t* transitioning to inactive ramet (IA) in *t+1* on whether ramet *i* in stage *x* in *t* survived in year *t*+1 (Equation 3). Thus, the model for annual transition probability of ramet *i* in stage *x* in year *t* to IA in *t+1* is given as:

$$logit[e(x_{i,t} \rightarrow y_{i,t+1} | surv_{i,x,t+1} )] = \beta_0 + (1|Year) \quad (3)$$

$$x_{i,t} \rightarrow y_{i,t+1} \sim Bernoulli[e(x_{i,t} \rightarrow y_{i,t+1})]$$

We conditioned the probability of ramet *i* in stage y in year *t* transitioning to either flowering single rosette (SR) or flowering multiple rosette (MR) stage in *t+1* on whether ramet *i* in stage *x* in *t* survived in year *t+1* and was not inactive (IA) in year *t* (Equation 4). Thus, the model for the annual transition probability of a ramet *i* in stage *y* in year *t* to SF stage, or to MF stage, in *t+1* is given as:

$$logit[e(x_{i,t} \rightarrow y_{i,t+1} | surv_{i,x,t+1}, not\ IA_t )] = \beta_0 + (1|Year) \quad (4)$$

$$x_{i,t} \rightarrow y_{i,t+1} \sim Bernoulli[e(x_{i,t} \rightarrow y_{i,t+1})]$$

***Recruitment.*** Ramets were recruited either as vegetative sprouts from underground taproots, linked it to an unknown single rosette (SR) or multiple rosette (MR) ramet (*x*) in year *t,* or as seedlings. Ramets recruited as sprouts from a taproot were seen for the first time as one of four stages: single rosette ($SR_{rec}$), multiple rosette ($MR_{rec}$), flowering single rosette ($SF_{rec}$), or flowering multiple rosette ($MF_{rec}$) in year *t+1*. We summed the annual number in each of these stages to get estimated total sprout recruits ($sprout_{rec}$). The number for each of these ramet stages (*y*) was estimated as the log of the expected number of sprout recruits in year *t+1* as a proportion of the total number of SR, or of MR, ramets in year *t;* this rule was used because Metcalf et al. (2009) found that the total number of ramets in a specific location directly affected the rate of establishment of new vegetative ramets. Thus, the model for annual sprout recruitment, calculated specifically for each ramet stage [single rosette ($SR_{rec}$), multiple rosette ($MR_{rec}$), flowering single rosette ($SF_{rec}$), and flowering multiple rosette ($MF_{rec}$)] is given as:

$$log[e(\frac{\sum sprout_{rec,y,t+1}}{\sum_{x=1}^{2} x_t})] = \beta_0 + (1|Year) \tag{5}$$

$$sprout_{rec,y,t+1} \sim poisson\ [e\left(\frac{\sum sprout_{rec,y,t+1}}{\sum_{x=1}^{2} x_t}\right)]$$

Ramets recruited as seedlings were identified in the field by size plus evidence of cotyledon leaves. Since no soil seed bank was found for *Cirsium* spp. at Arapaho (Potvin 1988), we assumed that all seedlings in year *t+1* germinated from seeds produced by flowering ramets (F) in year *t.* We estimated the annual number of recruited seedlings (Equation 6) as the sum of log of the expected number of seedlings in year *t+1* by each of the flowering ramet stages (SF, MF); the expected number in each stage for *t+1* was estimated as a proportion to the total number of flowering ramets in that stage, SF or MF, in year *t*. Thus, the model for annual seedling recruitment is given as:

$$log[e(\frac{\sum seedling_{rec,t+1}}{\sum_{f=1}^{2} f_t})] = \beta_0 + (1|Year) \tag{6}$$

$$seedling_{rec,t+1} \sim poisson\ [e\left(\frac{\sum seedling_{rec,t+1}}{\sum_{f=1}^{2} f_t}\right)]$$

### 3.0 DEVELOPMENT OF MATRIX POPULATION MODEL (MPM)

We synthesized the effect of annual variation in vital rates using population projection matrix models to predict annual population growth rates (Figure 1, main manuscript), with $n_{t+1} = A_t \times n_t$, where $n_t$ and $A_t$ are population vector and matrix (Equation 7) for year *t*, respectively. The matrix summarizes the transition probabilities from stage *x* in *t* (rows) to stage *y* in *t+1* (columns):

$$A_t = \begin{bmatrix} 0 & \delta_{sd} * Sd_{SR} & 0 & 0 & 0 & 0 \\ 0 & (\delta_{SR} * SR_{SR}) + SR_{rec} & (\delta_{SR} * SR_{SF}) + SF_{rec} & (\delta_{SR} * SR_{MR}) + MR_{rec} & (\delta_{SR} * SR_{MF}) + MR_{rec} & \delta_{SR} * SR_{IA} \\ SF_{sd} & 0 & 0 & 0 & 0 & 0 \\ 0 & (\delta_{MR} * MR_{SR}) + SR_{rec} & (\delta_{MR} * MR_{SF}) + SF_{rec} & (\delta_{MR} * MR_{MR}) + MR_{rec} & (\delta_{MR} * MR_{MF}) + MF_{rec} & \delta_{MR} * MR_{IA} \\ MF_{sd} & 0 & 0 & 0 & 0 & 0 \\ 0 & \delta_{IA} * IA_{SR} & \delta_{IA} * IA_{SF} & \delta_{IA} * IA_{MR} & 0 & \delta_{IA} * IA_{IA} \end{bmatrix} \quad (7)$$

The transition probabilities for each ramet stage in Equation 7 (Table S1) are defined as follows:

***Seedling, sd*** – Seedlings are recruited as ramets from the two flowering stages: either flowering single rosette ($SF_{sd}$) or flowering multiple rosette ($MF_{sd}$). All seedlings that survived their first year ($\delta_{sd}$) transitioned to ramets in the single rosette stage ($Sd_{SR}$).

***Single Rosette, SR*** – All ramets in the single rosette stage that survived a particular year ($\delta_{SR}$) either remained as a single rosette ($SR_{SR}$) or transitioned to one of four other stages: flowering single rosette ($SR_{SF}$), multiple rosette ($SR_{MR}$), flowering multiple rosette ($SR_{MF}$) or inactive ramet ($SR_{IA}$). A single rosette (SR) could also appear for the first time in *t* as a single rosette recruit ($SR_{rec}$) by vegetative reproduction from an unknown taproot.

***Multiple Rosette, MR*** – All ramets in the multiple rosette stage that survived a particular year ($\delta_{MR}$) could transition to one of four/five stages. It could: remain a multiple rosette ($MR_{MR}$), or became a single rosette ($MR_{SR}$) when all except one of its subrosettes died; or, it could become a flowering multiple rosette (*MR<sub>MF</sub>*), or a flowering single rosette ($MR_{SF}$); or, finally, it could become an inactive ramet ($MR_{IA}$). Note, since multiple rosettes varied in size, with up to seven sub-rosettes, although high numbers of subrosettes were uncommon; these ramets with multiple rosettes could lose or gain subrosettes and remain in the MR stage here.

***Inactive rosette, IA*** – All ramets that were inferred to be inactive, i.e., were missing for 1 or 2 years but seen again (within 3 years) had 100% survival by definition ($\delta_{IA}$). When inactive (IA), a ramet could either remain inactive for another year ($IA_{IA}$) or emerge after one year of inactivity. Ramets re-emerging at a tag were observed to appear in one of three stages: single rosette ($IA_{SR}$), flowering single rosette ($IA_{SF}$), or multiple rosette ($IA_{MR}$).

***Flowering rosettes: Flowering single rosette, SF, and flowering multiple rosette, MF*** – Each ramet of wavyleaf thistle that flowered died after flowering; however, it was associated with the recruitment of seedlings from the seeds produced. Ramets that flowered were observed transitioning from one of four stages: single rosette (SR); multiple rosette (MR); single flowering rosette sprout recruit ($SF_{rec}$) or multiple flowering rosette sprout recruit ($MF_{rec}$).

### 4.0 ASYMPTOTIC POPULATION GROWTH RATE

We calculated the dominant eigenvalue $\lambda_t$, the long-term asymptotic population growth rate estimated from year *t*, using the "popdemo" package in R (ver 4.4.1, R Core Team 2024). $\lambda_t$, is a good estimate of population viability if the population has reached the stable stage distribution (Tenhumberg, 2010). However, in some environments where populations are frequently perturbed away from the stable stage distribution, the short-term transient population growth rate is a better estimate of population viability (Caswell, 2001). Hence, we also calculated the transient growth rate, as:

$e^{\left(\frac{N_{t+1}-N_t}{N_t}\right)}$, where $N_t$= ramet population in year $t$ and $N_{t+1}$= ramet population in year $t+1$

### 5.0 WEATHER DATA

We downloaded monthly total rainfall and mean temperature data for each site from the PRISM Climate Group (Daly and Bryant, 2013); weather stations were at or near each site. We calculated the Standardized Precipitation Evapotranspiration Index (SPEI) (Vicente-Serrano et al., 2010a), using the SPEI package in R (4.4.1; R Core Team 2024). SPEI is a multi-scalar drought index that evaluates onset, duration, and severity of drought events relative to normal conditions within a specific geographic region (Vicente-Serrano et al., 2010a).

To calculate SPEI, we first calculated the potential evapotranspiration (PET) (Thornthwaite, 1948), using the monthly mean temperature and latitude of each study site (using the "thornthwaite" function in R). Then, we calculated SPEI values by quantifying the difference between monthly precipitation (PPT) and potential evapotranspiration (PET), aggregated over various time scales (measured in months) as follows (Vicente-Serrano et al., 2010b):

$$D_n^k = \sum_{i=0}^{k-1}(PPT_{n-i} - PET_{n-i}),\ n \geq k \qquad (8)$$

where *k* (months) is the time scale of the aggregation and *n* (years) is the length of the time series.

In all statistical analyses of the relationships between weather variables and demographic parameters, we used monthly data: mean monthly temperature anomaly, total monthly precipitation anomaly, and one-month SPEI.

For data visualization of weather variables, we used mean annual values (i.e., averaging the monthly data over every 12 months; from January – December of each year) to produce clearer patterns, including in SPEI where we plot 12-month SPEI.

TABLE S1: Description of vital rates and the associated parameters

| *Vital rates* | *Parameter* |
|---|---|
| ***Survival (*$\delta$*)*** | |
| Seedling (Sd) | $\delta_{sd}$ |
| Multiple Rosette-ramet (MR) | $\delta_{MR}$ |
| Single Rosette-ramet (SR) | $\delta_{SR}$ |
| Inactive Rosette-ramet (IR) | $\delta_{IA}$ |
| ***Sprout recruitment (rec) of*** | |
| Single Rosette-ramet (SR) | $SR_{rec}$ |
| Multiple Rosette-ramet (MR) | $MR_{rec}$ |
| Flowering Single Rosette-ramet (SF) | $SF_{rec}$ |
| Flowering Multiple Rosette-ramet (MF) | $MF_{rec}$ |
| ***Seedling (Sd) recruitment from*** | |
| Flowering Single Rosette-ramet (SF) | $SF \rightarrow Sd$ |
| Flowering Multiple Rosette-ramet (MF) | $MF \rightarrow Sd$ |
| ***Transition from Seedling (Sd) to*** | |
| Single Rosette-ramet (SR) | $Sd \rightarrow SR$ |
| ***Transition from Single Rosette (SR) to*** | |
| Single Rosette- ramet (SR) | $SR \rightarrow SR$ |
| Flowering Single Rosette- ramet (SF) | $SR \rightarrow SF$ |
| Multiple Rosette- ramet (MR) | $SR \rightarrow MR$ |

| | |
|---|---|
| Flowering Multiple Rosette- ramet (MF) | $SR \rightarrow MF$ |
| Inactive Rosette-ramet (IA) | $SR \rightarrow IA$ |
| ***Transition from Multiple Rosette (MR) to*** | |
| Single Rosette-ramet (SR) | $MR \rightarrow SR$ |
| Flowering Single Rosette-ramet (SF) | $MR \rightarrow SF$ |
| Multiple Rosette-ramet (MR) | $MR \rightarrow MR$ |
| Flowering Multiple Rosette-ramet (MF) | $MR \rightarrow MF$ |
| ***Transition from Inactive Rosette (IA) to*** | |
| Inactive Rosette-ramet (IA) | $IA \rightarrow IA$ |
| Single Rosette-ramet (SR) | $IA \rightarrow SR$ |
| Flowering Single Rosette-ramet (SF) | $IA \rightarrow SF$ |
| Multiple Rosette-ramet (MR) | $IA \rightarrow MR$ |
