## Appendix S2 for "Delayed effects of water-limiting conditions influence the ramet demography of a native iterocarpic thistle"

Table S1: Summary of annual vital rate estimates and asymptotic population growth rate of *C. undulatum* ramets at Arapaho.

| Year | $\delta_{IA}$ | IA→SF | IA→IA | IA→MR | IA→SR | $\delta_{SR}$ | SR→IA | SR→SF | SR→MF | SR→MR | SR→SR | $\delta_{MR}$ |
|---|---|---|---|---|---|---|---|---|---|---|---|---|
| 1991 | 1 | 0.05 | 0.02 | 0.05 | 0.95 | 0.65 | 0.07 | 0.17 | 0.00 | 0.02 | 0.97 | 0.77 |
| 1992 | 1 | 0.05 | 0.02 | 0.05 | 0.95 | 0.69 | 0.05 | 0.32 | 0.01 | 0.07 | 0.93 | 0.78 |
| 1993 | 1 | 0.05 | 0.02 | 0.05 | 0.95 | 0.72 | 0.01 | 0.44 | 0.00 | 0.05 | 0.95 | 0.68 |
| 1994 | 1 | 0.05 | 0.02 | 0.05 | 0.95 | 0.66 | 0.02 | 0.24 | 0.00 | 0.04 | 0.96 | 0.81 |
| 1995 | 1 | 0.05 | 0.02 | 0.05 | 0.95 | 0.60 | 0.01 | 0.15 | 0.00 | 0.03 | 0.97 | 0.85 |
| 1996 | 1 | 0.05 | 0.02 | 0.05 | 0.95 | 0.70 | 0.03 | 0.09 | 0.00 | 0.03 | 0.97 | 0.72 |
| 1997 | 1 | 0.05 | 0.02 | 0.05 | 0.95 | 0.65 | 0.04 | 0.10 | 0.00 | 0.02 | 0.97 | 0.74 |
| 1998 | 1 | 0.05 | 0.02 | 0.05 | 0.95 | 0.66 | 0.01 | 0.07 | 0.00 | 0.07 | 0.93 | 0.81 |
| 1999 | 1 | 0.05 | 0.02 | 0.05 | 0.95 | 0.64 | 0.01 | 0.12 | 0.00 | 0.04 | 0.96 | 0.61 |
| 2000 | 1 | 0.05 | 0.02 | 0.05 | 0.95 | 0.70 | 0.03 | 0.07 | 0.00 | 0.03 | 0.97 | 0.77 |
| 2001 | 1 | 0.05 | 0.02 | 0.05 | 0.95 | 0.43 | 0.09 | 0.28 | 0.01 | 0.03 | 0.97 | 0.59 |
| 2002 | 1 | 0.05 | 0.02 | 0.05 | 0.95 | 0.62 | 0.01 | 0.01 | 0.00 | 0.18 | 0.83 | 0.66 |
| 2003 | 1 | 0.05 | 0.02 | 0.05 | 0.95 | 0.49 | 0.05 | 0.12 | 0.00 | 0.02 | 0.98 | 0.50 |
| 2004 | 1 | 0.05 | 0.02 | 0.05 | 0.95 | 0.52 | 0.02 | 0.04 | 0.00 | 0.05 | 0.95 | 0.77 |
| 2005 | 1 | 0.05 | 0.02 | 0.05 | 0.95 | 0.71 | 0.07 | 0.11 | 0.00 | 0.03 | 0.97 | 0.69 |
| 2006 | 1 | 0.05 | 0.02 | 0.05 | 0.95 | 0.67 | 0.08 | 0.18 | 0.00 | 0.05 | 0.95 | 0.67 |
| 2007 | 1 | 0.05 | 0.02 | 0.05 | 0.95 | 0.74 | 0.06 | 0.24 | 0.01 | 0.04 | 0.96 | 0.81 |
| 2008 | 1 | 0.05 | 0.02 | 0.05 | 0.95 | 0.66 | 0.02 | 0.02 | 0.00 | 0.06 | 0.95 | 0.72 |
| 2009 | 1 | 0.05 | 0.02 | 0.05 | 0.95 | 0.89 | 0.00 | 0.06 | 0.00 | 0.02 | 0.98 | 0.87 |
| Year | MR→SF | MR→MF | MR→IA | MR→MR | MR→SR | $\delta_{seedling}$ | Seedling→SR | $MR_{rec}$ | $SR_{rec}$ | $SF_{rec}$ | $MF_{rec}$ | $Seedling_{rec}$ |
| 1991 | 0.12 | 0.06 | 0.06 | 0.20 | 0.72 | 0.69 | 1.00 | 0.01 | 0.30 | 0.01 | 0.00 | 0.70 |
| 1992 | 0.08 | 0.19 | 0.02 | 0.50 | 0.53 | 0.28 | 1.00 | 0.01 | 0.50 | 0.05 | 0.00 | 0.09 |
| 1993 | 0.07 | 0.27 | 0.02 | 0.20 | 0.72 | 0.69 | 1.00 | 0.04 | 1.72 | 0.13 | 0.02 | 1.50 |

| | | | | | | | | | | | | |
|---|---|---|---|---|---|---|---|---|---|---|---|---|
| 1994 | 0.08 | 0.05 | 0.02 | 0.36 | 0.63 | 0.45 | 1.00 | 0.04 | 0.61 | 0.02 | 0.00 | 1.43 |
| 1995 | 0.12 | 0.06 | 0.01 | 0.53 | 0.49 | 0.10 | 1.00 | 0.02 | 0.44 | 0.02 | 0.00 | 0.46 |
| 1996 | 0.08 | 0.22 | 0.02 | 0.30 | 0.68 | 0.60 | 1.00 | 0.02 | 0.50 | 0.01 | 0.00 | 0.31 |
| 1997 | 0.08 | 0.05 | 0.04 | 0.33 | 0.65 | 0.58 | 1.00 | 0.01 | 0.46 | 0.01 | 0.00 | 0.02 |
| 1998 | 0.11 | 0.12 | 0.02 | 0.25 | 0.72 | 0.69 | 1.00 | 0.02 | 0.41 | 0.01 | 0.00 | 0.37 |
| 1999 | 0.08 | 0.13 | 0.03 | 0.42 | 0.58 | 0.45 | 1.00 | 0.02 | 0.35 | 0.00 | 0.00 | 0.18 |
| 2000 | 0.05 | 0.05 | 0.01 | 0.56 | 0.46 | 0.39 | 1.00 | 0.01 | 0.28 | 0.01 | 0.00 | 0.07 |
| 2001 | 0.08 | 0.17 | 0.02 | 0.21 | 0.75 | 0.58 | 1.00 | 0.02 | 0.15 | 0.01 | 0.00 | 0.75 |
| 2002 | 0.07 | 0.05 | 0.02 | 0.29 | 0.68 | 0.34 | 1.00 | 0.08 | 0.68 | 0.00 | 0.00 | 0.02 |
| 2003 | 0.09 | 0.06 | 0.12 | 0.16 | 0.80 | 0.50 | 1.00 | 0.01 | 0.20 | 0.00 | 0.00 | 0.92 |
| 2004 | 0.07 | 0.06 | 0.02 | 0.25 | 0.71 | 0.44 | 1.00 | 0.02 | 0.73 | 0.01 | 0.00 | 0.68 |
| 2005 | 0.11 | 0.11 | 0.02 | 0.72 | 0.36 | 0.66 | 1.00 | 0.02 | 0.82 | 0.01 | 0.00 | 0.23 |
| 2006 | 0.09 | 0.19 | 0.05 | 0.15 | 0.78 | 0.31 | 1.00 | 0.02 | 0.31 | 0.01 | 0.00 | 0.03 |
| 2007 | 0.06 | 0.46 | 0.07 | 0.37 | 0.62 | 0.50 | 1.00 | 0.05 | 0.80 | 0.02 | 0.03 | 0.45 |
| 2008 | 0.06 | 0.04 | 0.02 | 0.33 | 0.66 | 0.50 | 1.00 | 0.04 | 1.24 | 0.01 | 0.00 | 0.38 |
| 2009 | 0.08 | 0.06 | 0.01 | 0.78 | 0.25 | 0.84 | 1.00 | 0.01 | 0.45 | 0.01 | 0.00 | 8.10 |

Sd = seedling; SR = juvenile single rosette, SF = flowering single rosette, MR = juvenile multiple rosette, MF = flowering multiple rosette; and, IA = inactive stage; (B) Life cycle diagram representing the observed demography of *C. undulatum* ramet stages. Additional symbols represent: for recruits: recruited as a seedling = $Seedling_{rec,}$ and, recruited as a vegetative sprout in stage: MR = $MR_{rec}$, MF = $MF_{rec}$, SR = $SR_{rec}$, and SF = $SF_{rec}$; and, for survivorship: $\delta_{sd}$ = seedling survival, $\delta_{SR}$ = SR survival, $\delta_{MR}$ = MR survival, $\delta_{IA}$ = IA survival. Arrows indicate transitions recorded among ramet stages.

Table S2: Summary of annual vital rate estimates and asymptotic population growth rate of *C. undulatum* ramets at Niobrara

| Year | $\delta_{IA}$ | IAt→SF | IA→IA | IA→MR | IA→SR | $\delta_{SR}$ | SR→IA | SR→SF | SR→MF | SR→MR | SR→SR | $\delta_{MR}$ |
|---|---|---|---|---|---|---|---|---|---|---|---|---|
| 1991 | 1.00 | 0.01 | 0.04 | 0.03 | 0.97 | 0.74 | 0.01 | 0.05 | 0.00 | 0.03 | 0.96 | 0.85 |
| 1992 | 1.00 | 0.01 | 0.03 | 0.03 | 0.97 | 0.68 | 0.04 | 0.05 | 0.00 | 0.05 | 0.95 | 0.72 |
| 1993 | 1.00 | 0.01 | 0.03 | 0.02 | 0.98 | 0.64 | 0.06 | 0.05 | 0.00 | 0.04 | 0.95 | 0.79 |
| 1994 | 1.00 | 0.01 | 0.10 | 0.02 | 0.98 | 0.77 | 0.04 | 0.03 | 0.00 | 0.04 | 0.95 | 0.88 |
| 1995 | 1.00 | 0.01 | 0.02 | 0.07 | 0.93 | 0.71 | 0.03 | 0.03 | 0.00 | 0.08 | 0.92 | 0.87 |
| 1996 | 1.00 | 0.01 | 0.03 | 0.02 | 0.98 | 0.73 | 0.01 | 0.05 | 0.00 | 0.03 | 0.96 | 0.74 |
| 1997 | 1.00 | 0.01 | 0.03 | 0.03 | 0.97 | 0.62 | 0.08 | 0.01 | 0.00 | 0.06 | 0.94 | 0.78 |
| 1998 | 1.00 | 0.01 | 0.02 | 0.02 | 0.98 | 0.64 | 0.03 | 0.04 | 0.00 | 0.02 | 0.97 | 0.82 |
| 1999 | 1.00 | 0.01 | 0.03 | 0.02 | 0.98 | 0.55 | 0.02 | 0.01 | 0.00 | 0.06 | 0.94 | 0.76 |
| 2000 | 1.00 | 0.01 | 0.03 | 0.03 | 0.97 | 0.58 | 0.05 | 0.02 | 0.00 | 0.05 | 0.95 | 0.83 |
| 2001 | 1.00 | 0.02 | 0.11 | 0.02 | 0.98 | 0.57 | 0.30 | 0.03 | 0.00 | 0.03 | 0.96 | 0.80 |
| 2002 | 1.00 | 0.01 | 0.01 | 0.03 | 0.97 | 0.72 | 0.05 | 0.04 | 0.00 | 0.08 | 0.93 | 0.81 |
| 2003 | 1.00 | 0.01 | 0.16 | 0.03 | 0.97 | 0.64 | 0.08 | 0.06 | 0.00 | 0.07 | 0.93 | 0.63 |
| 2004 | 1.00 | 0.01 | 0.02 | 0.02 | 0.98 | 0.48 | 0.01 | 0.03 | 0.00 | 0.05 | 0.95 | 0.58 |
| 2005 | 1.00 | 0.01 | 0.04 | 0.03 | 0.97 | 0.60 | 0.06 | 0.22 | 0.00 | 0.03 | 0.96 | 0.76 |
| 2006 | 1.00 | 0.01 | 0.03 | 0.02 | 0.98 | 0.77 | 0.02 | 0.03 | 0.00 | 0.03 | 0.96 | 0.82 |
| 2007 | 1.00 | 0.01 | 0.27 | 0.03 | 0.97 | 0.55 | 0.01 | 0.08 | 0.00 | 0.06 | 0.94 | 0.83 |
| 2008 | 1.00 | 0.01 | 0.27 | 0.03 | 0.97 | 0.61 | 0.01 | 0.02 | 0.00 | 0.04 | 0.95 | 0.59 |
| 2009 | 1.00 | 0.01 | 0.03 | 0.03 | 0.97 | 0.73 | 0.01 | 0.09 | 0.00 | 0.05 | 0.95 | 0.82 |

| Year | MR→SF | MR→MF | MR→IA | MR→MR | MR→SR | $\delta_{Seedling}$ | Seedling→SR | $MR_{rec}$ | $SR_{rec}$ | $SF_{rec}$ | $MF_{rec}$ | $Seedling_{rec}$ |
|---|---|---|---|---|---|---|---|---|---|---|---|---|
| 1991 | 0.10 | 0.01 | 0.03 | 0.24 | 0.74 | 0.47 | 1.00 | 0.01 | 0.29 | 0.00 | 0.00 | 3.20 |
| 1992 | 0.08 | 0.01 | 0.03 | 0.25 | 0.74 | 0.47 | 1.00 | 0.01 | 0.33 | 0.00 | 0.00 | 1.09 |
| 1993 | 0.12 | 0.01 | 0.03 | 0.23 | 0.74 | 0.47 | 1.00 | 0.01 | 0.29 | 0.00 | 0.00 | 1.09 |

| | | | | | | | | | | | | |
|---|---|---|---|---|---|---|---|---|---|---|---|---|
| 1994 | 0.15 | 0.01 | 0.03 | 0.24 | 0.74 | 0.47 | 1.00 | 0.01 | 0.37 | 0.00 | 0.00 | 1.46 |
| 1995 | 0.08 | 0.01 | 0.03 | 0.23 | 0.74 | 0.47 | 1.00 | 0.01 | 0.31 | 0.01 | 0.00 | 1.67 |
| 1996 | 0.09 | 0.00 | 0.03 | 0.25 | 0.74 | 0.47 | 1.00 | 0.01 | 0.22 | 0.00 | 0.00 | 1.50 |
| 1997 | 0.08 | 0.01 | 0.03 | 0.22 | 0.74 | 0.47 | 1.00 | 0.01 | 0.29 | 0.00 | 0.00 | 1.00 |
| 1998 | 0.07 | 0.00 | 0.03 | 0.22 | 0.74 | 0.47 | 1.00 | 0.01 | 0.22 | 0.00 | 0.00 | 0.99 |
| 1999 | 0.08 | 0.01 | 0.03 | 0.25 | 0.74 | 0.47 | 1.00 | 0.01 | 0.35 | 0.00 | 0.00 | 1.00 |
| 2000 | 0.10 | 0.01 | 0.03 | 0.25 | 0.74 | 0.47 | 1.00 | 0.01 | 0.15 | 0.00 | 0.00 | 0.55 |
| 2001 | 0.08 | 0.01 | 0.03 | 0.24 | 0.74 | 0.47 | 1.00 | 0.01 | 0.20 | 0.00 | 0.00 | 4.63 |
| 2002 | 0.09 | 0.01 | 0.03 | 0.24 | 0.74 | 0.47 | 1.00 | 0.01 | 0.56 | 0.00 | 0.00 | 0.29 |
| 2003 | 0.10 | 0.01 | 0.03 | 0.23 | 0.74 | 0.47 | 1.00 | 0.01 | 0.20 | 0.00 | 0.00 | 1.16 |
| 2004 | 0.08 | 0.01 | 0.03 | 0.24 | 0.74 | 0.47 | 1.00 | 0.01 | 0.27 | 0.00 | 0.00 | 0.38 |
| 2005 | 0.08 | 0.01 | 0.03 | 0.24 | 0.74 | 0.47 | 1.00 | 0.01 | 0.56 | 0.00 | 0.00 | 2.92 |
| 2006 | 0.08 | 0.11 | 0.03 | 0.24 | 0.74 | 0.47 | 1.00 | 0.01 | 0.64 | 0.00 | 0.00 | 0.21 |
| 2007 | 0.08 | 0.09 | 0.03 | 0.24 | 0.74 | 0.47 | 1.00 | 0.01 | 0.61 | 0.00 | 0.00 | 2.02 |
| 2008 | 0.08 | 0.01 | 0.03 | 0.24 | 0.74 | 0.47 | 1.00 | 0.01 | 0.66 | 0.00 | 0.00 | 0.68 |
| 2009 | 0.11 | 0.01 | 0.03 | 0.25 | 0.74 | 0.47 | 1.00 | 0.01 | 0.85 | 0.01 | 0.00 | 0.38 |

Sd = seedling; SR = juvenile single rosette, SF = flowering single rosette, MR = juvenile multiple rosette, MF = flowering multiple rosette; and, IA = inactive stage; (B) Life cycle diagram representing the observed demography of *C. undulatum* ramet stages. Additional symbols represent: for recruits: recruited as a seedling = $Seedling_{rec}$, and, recruited as a vegetative sprout in stage: MR = $MR_{rec}$, MF = $MF_{rec}$, SR = $SR_{rec}$, and SF = $SF_{rec}$; and, for survivorship: $\delta_{sd}$ = seedling survival, $\delta_{SR}$ = SR survival, $\delta_{MR}$ = MR survival, $\delta_{IA}$ = IA survival. Arrows indicate transitions recorded among ramet stages.

Table S3: Summary of all FLMs fitted to ramet vital rates and population growth at Arapaho. Bold highlighted rows are FLMs that were significant at $p<0.01$

| Vital Rate | model | df | logLik | AIC | BIC | deviance | df.residual | adj.$r^2$ | edf | F-statistic | p.value | Delta_AIC |
|---|---|---|---|---|---|---|---|---|---|---|---|---|
| Recruitment of Seedling | Mean Temperature | 2.83 | -32.68 | 73.01 | 76.63 | 34.70 | 16.17 | 0.14 | 0.83 | 0.35 | 0.04 | 9.56 |
| | **Total Precipitation** | **3.00** | **-30.03** | **68.07** | **71.85** | **26.26** | **16.00** | **0.34** | **1.00** | **0.95** | **0.00** | **4.62** |
| | **Drought Index, SPEI** | **3.21** | **-27.51** | **63.45** | **67.43** | **20.14** | **15.79** | **0.49** | **1.21** | **1.67** | **0.00** | **0.00** |
| Recruitment of Flowering Multiple Rosette | Mean Temperature | 2.00 | -30.10 | 66.19 | 69.02 | 26.43 | 17.00 | 0.04 | 0.00 | 0.00 | 0.30 | 0.91 |
| | Total Precipitation | 2.00 | -30.10 | 66.19 | 69.02 | 26.43 | 17.00 | 0.04 | 0.00 | 0.00 | 0.50 | 0.91 |
| | Drought Index, SPEI | 2.48 | -29.16 | 65.28 | 68.56 | 23.96 | 16.52 | 0.11 | 0.48 | 0.11 | 0.14 | 0.00 |
| Recruitment of Flowering Single Rosette | Mean Temperature | 7.47 | -14.07 | 45.09 | 53.08 | 4.89 | 11.53 | 0.51 | 5.47 | 1.55 | 0.06 | 11.21 |
| | Total Precipitation | 9.12 | -9.86 | 39.96 | 49.51 | 3.14 | 9.88 | 0.63 | 7.12 | 2.66 | 0.03 | 6.08 |
| | **Drought Index, SPEI** | **7.52** | **-8.41** | **33.87** | **41.92** | **2.70** | **11.48** | **0.73** | **5.52** | **3.98** | **0.00** | **0.00** |
| Recruitment of Single Rosette | Mean Temperature | 2.65 | -15.20 | 37.71 | 41.16 | 5.51 | 16.35 | 0.07 | 0.65 | 0.20 | 0.08 | 2.35 |

| | | | | | | | | | | | | |
|---|---|---|---|---|---|---|---|---|---|---|---|---|
| | Total Precipitation | 3.60 | -13.78 | 36.76 | 41.11 | 4.75 | 15.40 | 0.15 | 1.60 | 0.46 | 0.09 | 1.41 |
| | Drought Index, SPEI | 3.57 | -13.11 | 35.35 | 39.67 | 4.42 | 15.43 | 0.21 | 1.57 | 0.60 | 0.04 | 0.00 |
| Recruitment of Multiple Rosette | Mean Temperature | 2.00 | -14.69 | 35.38 | 38.22 | 5.22 | 17.00 | -0.05 | 0.00 | 0.00 | 0.55 | 7.08 |
| | Total Precipitation Anomaly | 7.55 | -6.34 | 29.77 | 37.85 | 2.17 | 11.45 | 0.35 | 5.55 | 1.35 | 0.09 | 1.47 |
| | Drought Index, SPEI | 8.71 | -4.44 | 28.30 | 37.48 | 1.78 | 10.29 | 0.41 | 6.71 | 1.66 | 0.09 | 0.00 |
| Survival of Seedling | Mean Temperature | 2.84 | -20.62 | 48.92 | 52.54 | 9.75 | 16.16 | 0.16 | 0.84 | 0.33 | 0.05 | 0.00 |
| | Total Precipitation | 2.26 | -22.55 | 51.62 | 54.69 | 11.95 | 16.74 | 0.00 | 0.26 | 0.04 | 0.21 | 2.70 |
| | Drought Index, SPEI | 2.31 | -22.45 | 51.52 | 54.65 | 11.82 | 16.69 | 0.01 | 0.31 | 0.05 | 0.20 | 2.61 |
| Multiple Rosette to Single Rosette | Mean Temperature | 7.27 | -10.12 | 36.78 | 44.59 | 3.23 | 11.73 | 0.32 | 5.27 | 1.09 | 0.13 | 4.94 |
| | **Total Precipitation Anomaly** | **3.05** | **-11.87** | **31.84** | **35.66** | **3.88** | **15.95** | **0.40** | **1.05** | **1.08** | **0.00** | **0.00** |
| | Drought Index, SPEI | 2.80 | -15.02 | 37.65 | 41.24 | 5.41 | 16.20 | 0.18 | 0.80 | 0.35 | 0.04 | 5.81 |
| Multiple Rosette to Multiple Rosette | Mean Temperature | 7.12 | -15.27 | 46.78 | 54.45 | 5.55 | 11.88 | 0.31 | 5.12 | 1.04 | 0.14 | 3.52 |

| | | | | | | | | | | | | |
|---|---|---|---|---|---|---|---|---|---|---|---|---|
| | Total Precipitation | 2.99 | -17.64 | 43.26 | 47.03 | 7.13 | 16.01 | 0.34 | 0.99 | 0.84 | 0.01 | 0.00 |
| | Drought Index, SPEI | 2.75 | -20.24 | 47.97 | 51.51 | 9.36 | 16.25 | 0.15 | 0.75 | 0.29 | 0.05 | 4.71 |
| Multiple Rosette to Inactive | Mean Temperature | 2.00 | -18.66 | 43.33 | 46.16 | 7.93 | 17.00 | -0.05 | 0.00 | 0.00 | 0.59 | 0.80 |
| | Total Precipitation | 3.56 | -16.70 | 42.53 | 46.84 | 6.45 | 15.44 | 0.06 | 1.56 | 0.26 | 0.22 | 0.00 |
| | Drought Index, SPEI | 2.00 | -18.66 | 43.33 | 46.16 | 7.93 | 17.00 | -0.05 | 0.00 | 0.00 | 0.54 | 0.80 |
| Multiple Rosette to Flowering Multiple Rosette | Mean Temperature | 4.57 | -18.70 | 48.56 | 53.82 | 7.97 | 14.43 | 0.21 | 2.57 | 0.71 | 0.08 | 0.00 |
| | Total Precipitation | 2.00 | -23.04 | 52.08 | 54.91 | 12.58 | 17.00 | -0.06 | 0.00 | 0.00 | 0.84 | 3.52 |
| | Drought Index, SPEI | 2.00 | -23.04 | 52.08 | 54.91 | 12.58 | 17.00 | -0.06 | 0.00 | 0.00 | 0.44 | 3.52 |
| Multiple Rosette to Flowering Single Rosette | Mean Temperature | 2.00 | 1.50 | 2.99 | 5.82 | 0.95 | 17.00 | 0.05 | 0.00 | 0.00 | 0.89 | 5.51 |
| | Total Precipitation | 6.87 | 8.88 | -2.02 | 5.42 | 0.44 | 12.13 | 0.39 | 4.87 | 1.16 | 0.09 | 0.50 |
| | Drought Index, SPEI | 7.75 | 10.01 | -2.52 | 5.75 | 0.39 | 11.25 | 0.41 | 5.75 | 1.29 | 0.11 | 0.00 |
| Survival of Multiple Rosette | Mean Temperature | 2.00 | -12.65 | 31.31 | 34.14 | 4.21 | 17.00 | -0.05 | 0.00 | 0.00 | 0.59 | 3.92 |

| | | | | | | | | | | | | |
|---|---|---|---|---|---|---|---|---|---|---|---|---|
| | Total Precipitation Anomaly | 2.84 | -9.85 | 27.39 | 31.02 | 3.14 | 16.16 | 0.17 | 0.84 | 0.43 | 0.03 | 0.00 |
| | Drought Index, SPEI | 2.76 | -10.60 | 28.72 | 32.27 | 3.40 | 16.24 | 0.11 | 0.76 | 0.28 | 0.06 | 1.33 |
| Single Rosette to Single Rosette | Mean Temperature | 2.00 | -14.18 | 34.36 | 37.19 | 4.95 | 17.00 | -0.06 | 0.00 | 0.00 | 0.51 | 2.22 |
| | Total Precipitation | 2.72 | -12.35 | 32.14 | 35.66 | 4.08 | 16.28 | 0.09 | 0.72 | 0.25 | 0.07 | 0.00 |
| | Drought Index, SPEI | 2.60 | -12.89 | 32.99 | 36.39 | 4.32 | 16.40 | 0.04 | 0.60 | 0.16 | 0.11 | 0.85 |
| Single Rosette to Multiple Rosette | Mean Temperature | 2.00 | -16.96 | 39.92 | 42.75 | 6.63 | 17.00 | -0.06 | 0.00 | 0.00 | 0.53 | 1.86 |
| | Total Precipitation | 2.70 | -15.33 | 38.06 | 41.55 | 5.59 | 16.30 | 0.07 | 0.70 | 0.22 | 0.09 | 0.00 |
| | Drought Index, SPEI | 2.52 | -15.95 | 38.94 | 42.27 | 5.96 | 16.48 | 0.02 | 0.52 | 0.12 | 0.14 | 0.89 |
| **Single Rosette to Flowering Multiple Rosette** | **Mean Temperature Anomaly** | **7.41** | **-2.71** | **22.23** | **30.17** | **1.48** | **11.59** | **0.67** | **5.41** | **3.31** | **0.00** | **0.00** |
| | Total Precipitation | 6.13 | -10.65 | 35.57 | 42.30 | 3.41 | 12.87 | 0.31 | 4.13 | 0.86 | 0.13 | 13.34 |
| | Drought Index, SPEI | 2.06 | -16.34 | 38.81 | 41.70 | 6.21 | 16.94 | 0.05 | 0.06 | 0.01 | 0.35 | 16.58 |
| Single Rosette to Flowering Single Rosette | Mean Temperature | 5.31 | -20.19 | 53.00 | 58.96 | 9.31 | 13.69 | 0.44 | 3.31 | 0.92 | 0.07 | 0.00 |

| | | | | | | | | | | | | |
|---|---|---|---|---|---|---|---|---|---|---|---|---|
| | Total Precipitation | 2.20 | -25.50 | 57.39 | 60.40 | 16.29 | 16.80 | 0.20 | 0.20 | 0.03 | 0.23 | 4.39 |
| | Drought Index, SPEI | 4.87 | -20.65 | 53.04 | 58.59 | 9.78 | 14.13 | 0.43 | 2.87 | 0.90 | 0.05 | 0.04 |
| Single Rosette to Inactive | Mean Temperature | 4.99 | -18.38 | 48.75 | 54.41 | 7.70 | 14.01 | 0.33 | 2.99 | 1.11 | 0.03 | 0.00 |
| | Total Precipitation | 2.60 | -23.20 | 53.60 | 57.00 | 12.78 | 16.40 | 0.05 | 0.60 | 0.17 | 0.11 | 4.85 |
| | Drought Index, SPEI | 2.48 | -23.62 | 54.21 | 57.50 | 13.37 | 16.52 | 0.01 | 0.48 | 0.10 | 0.15 | 5.46 |
| Survival of Single Rosette | Mean Temperature | 2.00 | -12.39 | 30.79 | 33.62 | 4.10 | 17.00 | -0.02 | 0.00 | 0.00 | 0.42 | 10.80 |
| | **Total Precipitation** | **3.21** | **-5.78** | **19.98** | **23.96** | **2.04** | **15.79** | **0.45** | **1.21** | **1.39** | **0.00** | **0.00** |
| | **Drought Index, SPEI** | **3.00** | **-8.67** | **25.33** | **29.10** | **2.77** | **16.00** | **0.27** | **1.00** | **0.61** | **0.01** | **5.34** |
| Lambda | Mean Temperature | 2.83 | 0.54 | 6.59 | 10.21 | 1.05 | 16.17 | 0.17 | 0.83 | 0.38 | 0.03 | 6.14 |
| | **Total Precipitation** | **3.74** | **3.45** | **2.58** | **7.05** | **0.77** | **15.26** | **0.35** | **1.74** | **1.06** | **0.01** | **2.13** |
| | **Drought Index, SPEI** | **3.73** | **4.51** | **0.45** | **4.92** | **0.69** | **15.27** | **0.42** | **1.73** | **1.35** | **0.00** | **0.00** |

Table S4: Summary of all FLMs fitted to ramet vital rates and population growth at Niobrara. Bold highlighted rows are FLMs that were significant at *p*<0.01

| Vital Rate | model | df | logLik | AIC | BIC | deviance | df.residual | adj.$r^2$ | edf | statistic | p.value | delta AIC |
|---|---|---|---|---|---|---|---|---|---|---|---|---|
| Recruitment of Seedling | Mean Temperature | 1.00 | -21.14 | 46.29 | 48.18 | 10.30 | 18.00 | 0.16 | 0.00 | 0.00 | 0.97 | 11.18 |
| | **Total Precipitation** | **3.73** | **-13.33** | **36.12** | **40.59** | **4.52** | **15.27** | **0.57** | **1.73** | **1.77** | **0.00** | **1.01** |
| | **Drought Index, SPEI** | **3.84** | **-12.72** | **35.11** | **39.68** | **4.24** | **15.16** | **0.59** | **1.84** | **1.95** | **0.00** | **0.00** |
| Recruitment of Flowering Single Rosette | Mean Temperature | 2.00 | -18.65 | 43.30 | 46.13 | 7.92 | 17.00 | 0.04 | 0.00 | 0.00 | 0.56 | 0.00 |
| | Total Precipitation | 2.00 | -18.65 | 43.30 | 46.13 | 7.92 | 17.00 | 0.04 | 0.00 | 0.00 | 1.00 | 0.00 |
| | Drought Index, SPEI | 2.00 | -18.65 | 43.30 | 46.13 | 7.92 | 17.00 | 0.04 | 0.00 | 0.00 | 0.59 | 0.00 |
| Recruitment of Single Rosette | Mean Temperature | 2.00 | -9.53 | 25.05 | 27.89 | 3.03 | 17.00 | 0.26 | 0.00 | 0.00 | 0.79 | 4.37 |
| | Total Precipitation | 2.00 | -9.53 | 25.05 | 27.89 | 3.03 | 17.00 | 0.26 | 0.00 | 0.00 | 0.35 | 4.37 |
| | Drought Index, SPEI | 3.71 | -5.63 | 20.69 | 25.14 | 2.01 | 15.29 | 0.45 | 1.71 | 0.63 | 0.05 | 0.00 |
| Recruitment of Multiple Rosette | Mean Temperature | 2.00 | 3.83 | -1.66 | 1.17 | 0.74 | 17.00 | -0.06 | 0.00 | 0.00 | 0.30 | 0.34 |
| | Total Precipitation | 2.28 | 4.28 | -2.01 | 1.09 | 0.71 | 16.72 | -0.02 | 0.28 | 0.05 | 0.21 | 0.00 |

| | | | | | | | | | | | | |
|---|---|---|---|---|---|---|---|---|---|---|---|---|
| | Drought Index, SPEI | 2.00 | 3.83 | -1.66 | 1.17 | 0.74 | 17.00 | -0.06 | 0.00 | 0.00 | 0.67 | 0.34 |
| Multiple Rosette to Multiple Rosette | Mean Temperature | 2.61 | 33.36 | -59.49 | -56.08 | 0.03 | 16.39 | 0.05 | 0.61 | 0.14 | 0.13 | 0.00 |
| | Total Precipitation | 2.00 | 32.17 | -58.34 | -55.51 | 0.04 | 17.00 | -0.04 | 0.00 | 0.00 | 0.55 | 1.15 |
| | Drought Index, SPEI | 2.00 | 32.17 | -58.34 | -55.51 | 0.04 | 17.00 | -0.04 | 0.00 | 0.00 | 0.75 | 1.15 |
| Multiple Rosette to Flowering Multiple Rosette | Mean Temperature | 4.79 | -20.22 | 52.01 | 57.48 | 9.34 | 14.21 | 0.28 | 2.79 | 0.48 | 0.20 | 0.00 |
| | Total Precipitation | 2.00 | -23.54 | 53.09 | 55.92 | 13.26 | 17.00 | 0.14 | 0.00 | 0.00 | 0.94 | 1.07 |
| | Drought Index, SPEI | 2.00 | -23.54 | 53.09 | 55.92 | 13.26 | 17.00 | 0.14 | 0.00 | 0.00 | 0.77 | 1.07 |
| Multiple Rosette to Flowering Single Rosette | Mean Temperature | 4.75 | 6.63 | -1.77 | 3.66 | 0.55 | 14.25 | 0.11 | 2.75 | 0.39 | 0.28 | 0.00 |
| | Total Precipitation | 2.00 | 3.68 | -1.36 | 1.47 | 0.75 | 17.00 | -0.02 | 0.00 | 0.00 | 0.55 | 0.40 |
| | Drought Index, SPEI | 2.00 | 3.68 | -1.36 | 1.47 | 0.75 | 17.00 | -0.02 | 0.00 | 0.00 | 0.89 | 0.40 |
| Survival of Multiple Rosette | Mean Temperature | 2.00 | -10.85 | 27.70 | 30.54 | 3.49 | 17.00 | 0.07 | 0.00 | 0.00 | 0.82 | 0.00 |

| | | | | | | | | | | | | |
|---|---|---|---|---|---|---|---|---|---|---|---|---|
| | Total Precipitation | 2.00 | -10.85 | 27.70 | 30.54 | 3.49 | 17.00 | 0.07 | 0.00 | 0.00 | 0.70 | 0.00 |
| | Drought Index, SPEI | 2.00 | -10.85 | 27.70 | 30.54 | 3.49 | 17.00 | 0.07 | 0.00 | 0.00 | 0.89 | 0.00 |
| Single Rosette to Single Rosette | Mean Temperature | 2.00 | 1.55 | 2.91 | 5.74 | 0.95 | 17.00 | -0.06 | 0.00 | 0.00 | 0.91 | 0.00 |
| | Total Precipitation | 2.00 | 1.55 | 2.91 | 5.74 | 0.95 | 17.00 | -0.06 | 0.00 | 0.00 | 0.70 | 0.00 |
| | Drought Index, SPEI | 2.00 | 1.55 | 2.91 | 5.74 | 0.95 | 17.00 | -0.06 | 0.00 | 0.00 | 0.85 | 0.00 |
| Single Rosette to Multiple Rosette | Mean Temperature | 2.00 | -7.34 | 20.68 | 23.52 | 2.41 | 17.00 | -0.06 | 0.00 | 0.00 | 0.90 | 0.00 |
| | Total Precipitation | 2.00 | -7.34 | 20.68 | 23.52 | 2.41 | 17.00 | -0.06 | 0.00 | 0.00 | 0.63 | 0.00 |
| | Drought Index, SPEI | 2.00 | -7.34 | 20.68 | 23.52 | 2.41 | 17.00 | -0.06 | 0.00 | 0.00 | 0.82 | 0.00 |
| Single Rosette to Flowering Single Rosette | Mean Temperature | 2.00 | -20.71 | 47.42 | 50.26 | 9.84 | 17.00 | 0.00 | 0.00 | 0.00 | 0.90 | 0.00 |
| | Total Precipitation | 2.01 | -20.70 | 47.42 | 50.26 | 9.83 | 16.99 | 0.00 | 0.01 | 0.00 | 0.34 | 0.00 |
| | Drought Index, SPEI | 2.00 | -20.71 | 47.42 | 50.26 | 9.84 | 17.00 | 0.00 | 0.00 | 0.00 | 0.31 | 0.00 |

| | | | | | | | | | | | | |
|---|---|---|---|---|---|---|---|---|---|---|---|---|
| Single Rosette to Inactive | Mean Temperature | 2.00 | -25.19 | 56.37 | 59.20 | 15.76 | 17.00 | -0.03 | 0.00 | 0.00 | 0.45 | 3.93 |
| | Total Precipitation | 3.67 | -21.55 | 52.44 | 56.86 | 10.75 | 15.33 | 0.22 | 1.67 | 0.58 | 0.06 | 0.00 |
| | Drought Index, SPEI | 3.30 | -23.32 | 55.25 | 59.32 | 12.95 | 15.70 | 0.08 | 1.30 | 0.24 | 0.19 | 2.81 |
| Survival of Single Rosette | Mean Temperature | 2.00 | -6.46 | 18.93 | 21.76 | 2.20 | 17.00 | 0.04 | 0.00 | 0.00 | 0.67 | 1.46 |
| | Total Precipitation | 2.00 | -6.46 | 18.93 | 21.76 | 2.20 | 17.00 | 0.04 | 0.00 | 0.00 | 0.46 | 1.46 |
| | Drought Index, SPEI | 3.60 | -4.14 | 17.47 | 21.81 | 1.72 | 15.40 | 0.17 | 1.60 | 0.31 | 0.17 | 0.00 |
| Inactive to Single Rosette | Mean Temperature | 4.74 | 1.89 | 7.70 | 13.12 | 0.91 | 14.26 | 0.05 | 0.00 | 0.00 | 0.94 | 0.74 |
| | Total Precipitation | 2.24 | -0.24 | 6.96 | 10.02 | 1.14 | 16.76 | -0.01 | 0.24 | 0.04 | 0.22 | 0.00 |
| | Drought Index, SPEI | 2.00 | -0.62 | 7.24 | 10.08 | 1.19 | 17.00 | -0.04 | 0.00 | 0.00 | 0.42 | 0.28 |
| Inactive to Multiple Rosette | Mean Temperature | 4.74 | 1.89 | 7.70 | 13.12 | 0.91 | 14.26 | 0.05 | 0.00 | 0.00 | 0.94 | 0.74 |
| | Total Precipitation | 2.24 | -0.24 | 6.96 | 10.02 | 1.14 | 16.76 | -0.01 | 0.24 | 0.04 | 0.22 | 0.00 |
| | Drought Index, SPEI | 2.00 | -0.62 | 7.24 | 10.08 | 1.19 | 17.00 | -0.04 | 0.00 | 0.00 | 0.42 | 0.28 |
| Inactive to Inactive | Mean Temperature | 5.70 | -20.02 | 53.44 | 59.77 | 9.15 | 13.30 | 0.24 | 3.70 | 0.55 | 0.26 | 4.77 |

| | | | | | | | | | | | | |
|---|---|---|---|---|---|---|---|---|---|---|---|---|
| | Total Precipitation | 2.34 | -23.53 | 53.74 | 56.89 | 13.25 | 16.66 | 0.13 | 0.34 | 0.06 | 0.19 | 5.07 |
| | Drought Index, SPEI | 7.69 | -15.64 | 48.67 | 56.88 | 5.77 | 11.31 | 0.44 | 5.69 | 1.34 | 0.10 | 0.00 |
| Inactive to Flowering Single Rosette | Mean Temperature | 3.59 | 12.28 | -15.37 | -11.03 | 0.31 | 15.41 | 0.08 | 1.59 | 0.30 | 0.18 | 0.00 |
| | Total Precipitation | 2.00 | 10.05 | -14.10 | -11.27 | 0.39 | 17.00 | -0.05 | 0.00 | 0.00 | 0.86 | 1.26 |
| | Drought Index, SPEI | 5.06 | 13.59 | -15.06 | -9.34 | 0.27 | 13.94 | 0.11 | 3.06 | 0.49 | 0.22 | 0.31 |
| Lambda | Mean Temperature | 1.00 | 6.86 | -9.73 | -7.84 | 0.54 | 18.00 | 0.15 | 0.00 | 0.00 | 0.40 | 0.35 |
| | Total Precipitation | 1.00 | 6.86 | -9.73 | -7.84 | 0.54 | 18.00 | 0.15 | 0.00 | 0.00 | 0.33 | 0.35 |
| | Drought Index, SPEI | 3.44 | 9.49 | -10.08 | -5.89 | 0.41 | 15.56 | 0.22 | 1.44 | 0.30 | 0.15 | 0.00 |

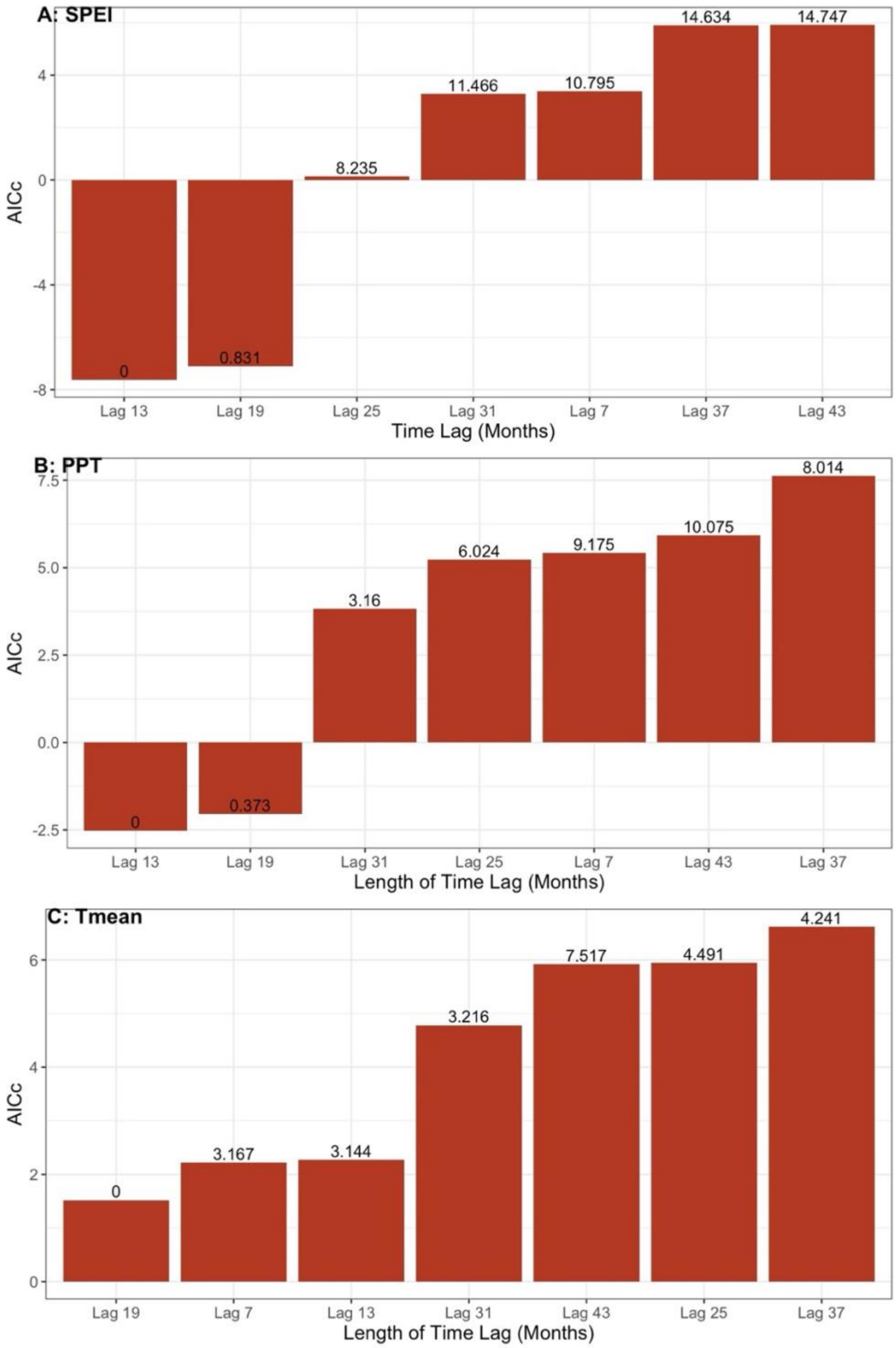


FIGURE S1: Model selection results for time lags of 7, 13, 19, 25, 31, 37, and 43 months, evaluated using the corrected Akaike Information Criterion (AICc). The values on the bars are the calculated AICc.

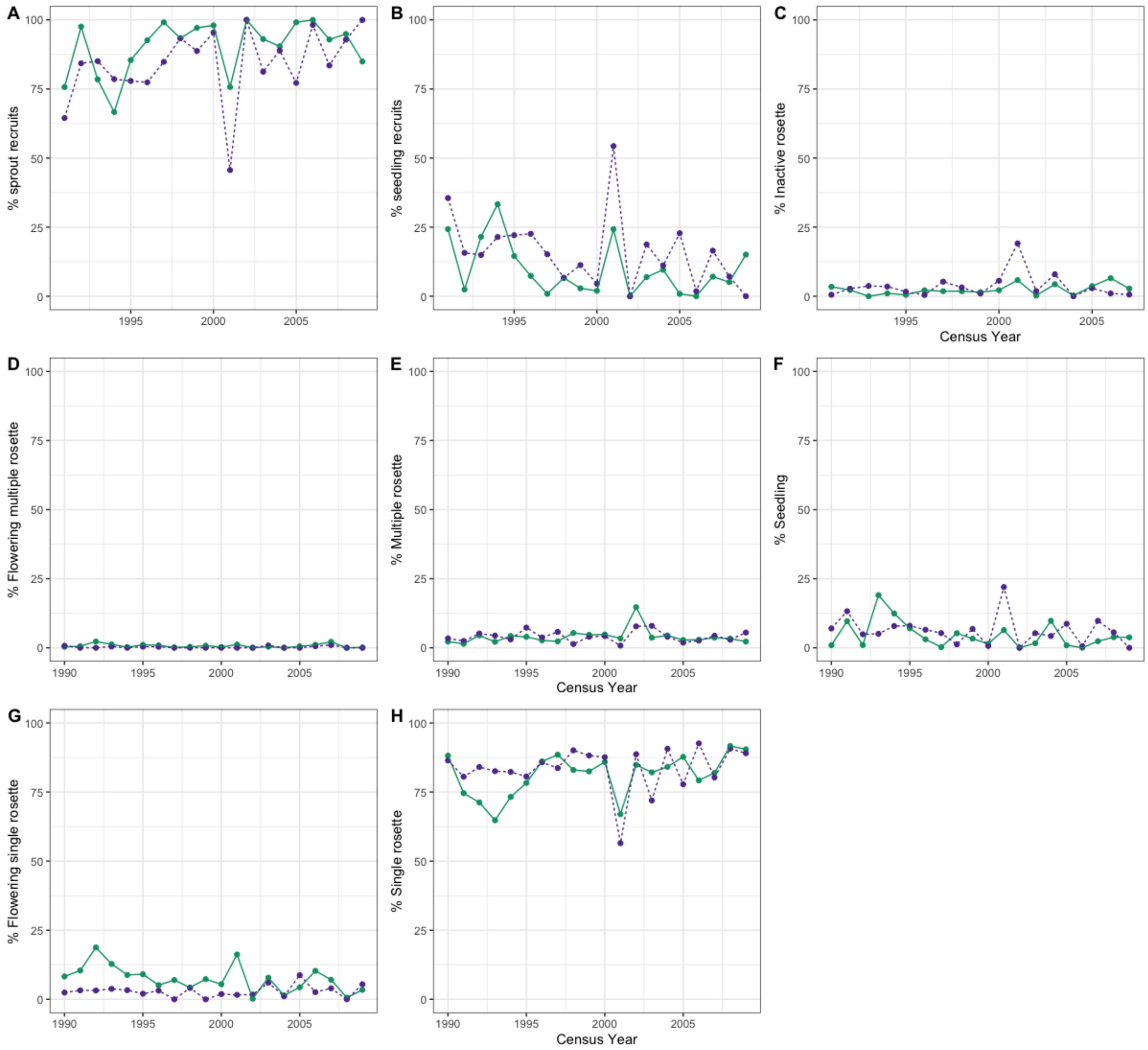


FIGURE S2. The proportion of ramets in each stage out of the total ramets observed at Arapaho (solid line) versus at Niobrara (broken line). Panels A & B were calculated based only on the ramets of known age, those that appeared for the first time during the study. Panels C-H) illustrate the proportion of ramets in each of the six transition stages calculated from the total ramets observed. Panel (A) vs Panel (B) compares the proportion of sprout recruits to the proportion of seedling recruits, respectively: Panel A and Panel B show that recruitment was predominantly by vegetative sprouts (Panel B) rather than by seedling establishment (Panel A). Panels C, D, E, F, G, and H compare the proportions among the six ramet stages, inactive, Flowering multiple rosette, Multiple rosette, Seedling, Flowering single rosette, Single rosette: Panels C, D, E, F, G, and H show that Single rosette is the most predominant stage among the six ramet stages.

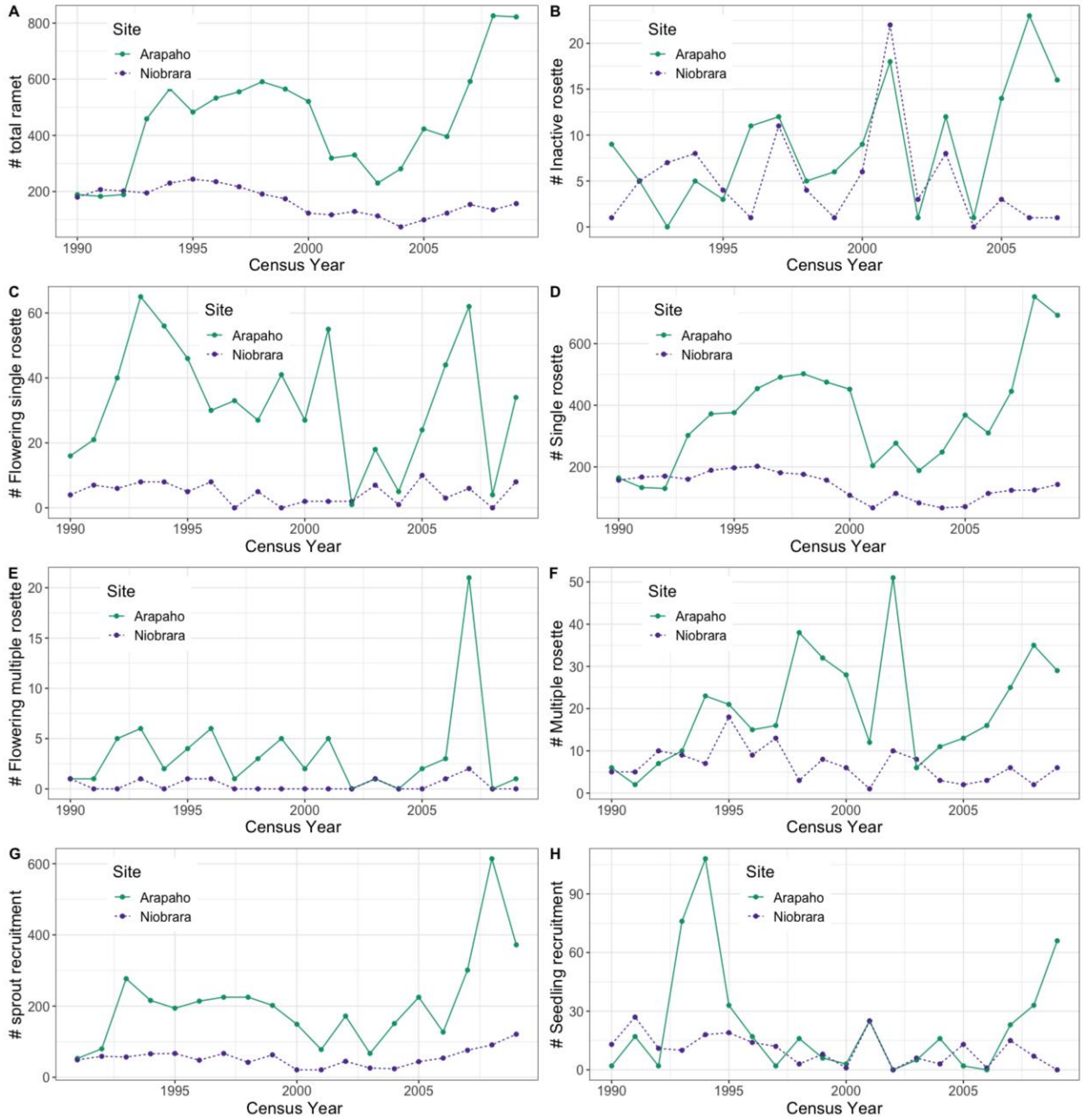


FIGURE S3: Comparing the dynamics of (A) the total number of ramets, (B) recruitment of sprouts, (C) recruitment of seedlings, (D-H) the number of ramets per stage. To manage the prairie, plots were hayed in late July after the population census in a four-year rotation cycle at Arapaho (1989, 1993, 1997, 2001, 2005) and lightly grazed (≤150 cow-calf pairs) early (May) and late (September) in the growing season at Niobrara.

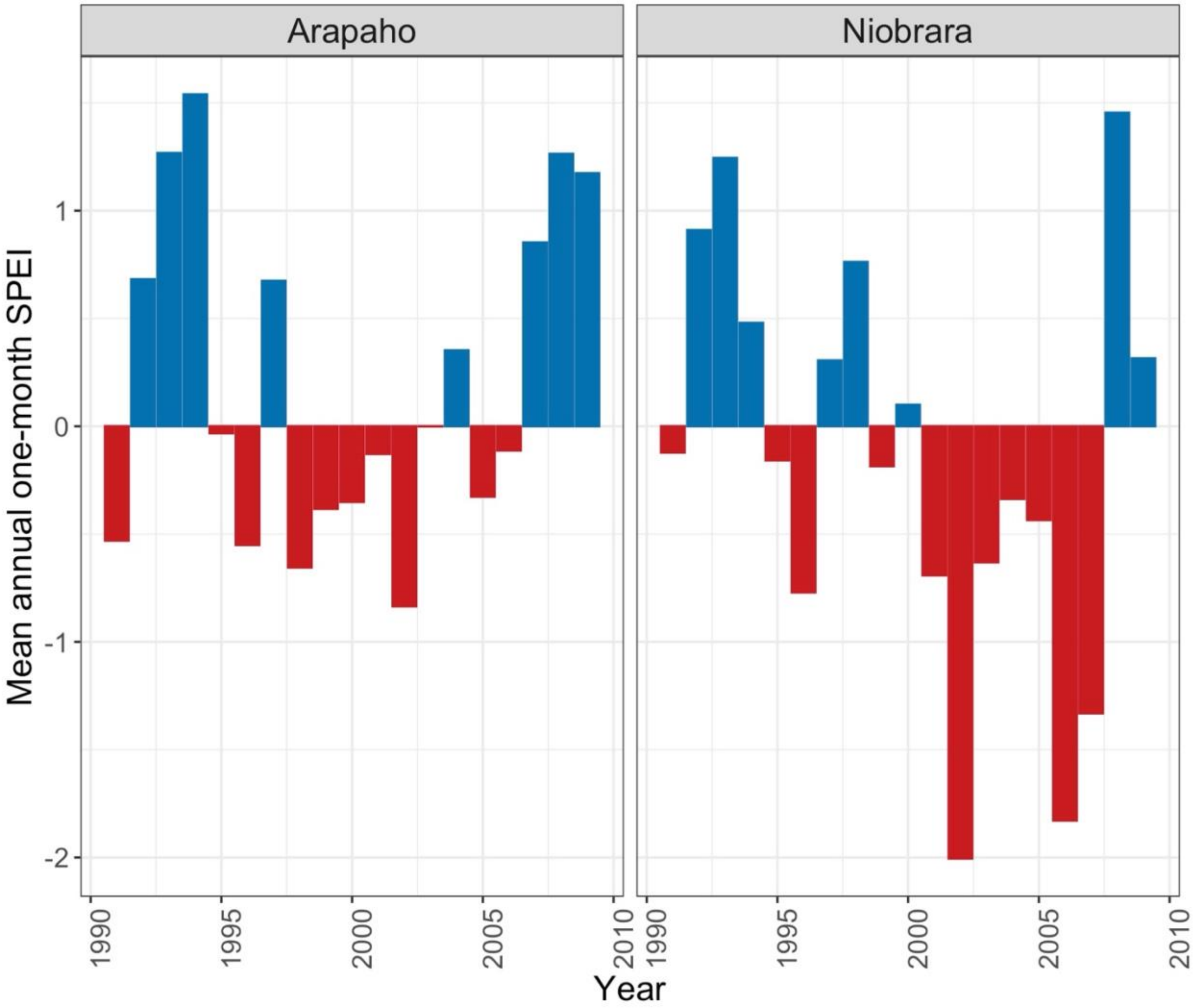


FIGURE S4: Annual average SPEI from June of the previous year (t–1) to July of the census year (t): Blue bars = wet years, red bars = dry years.

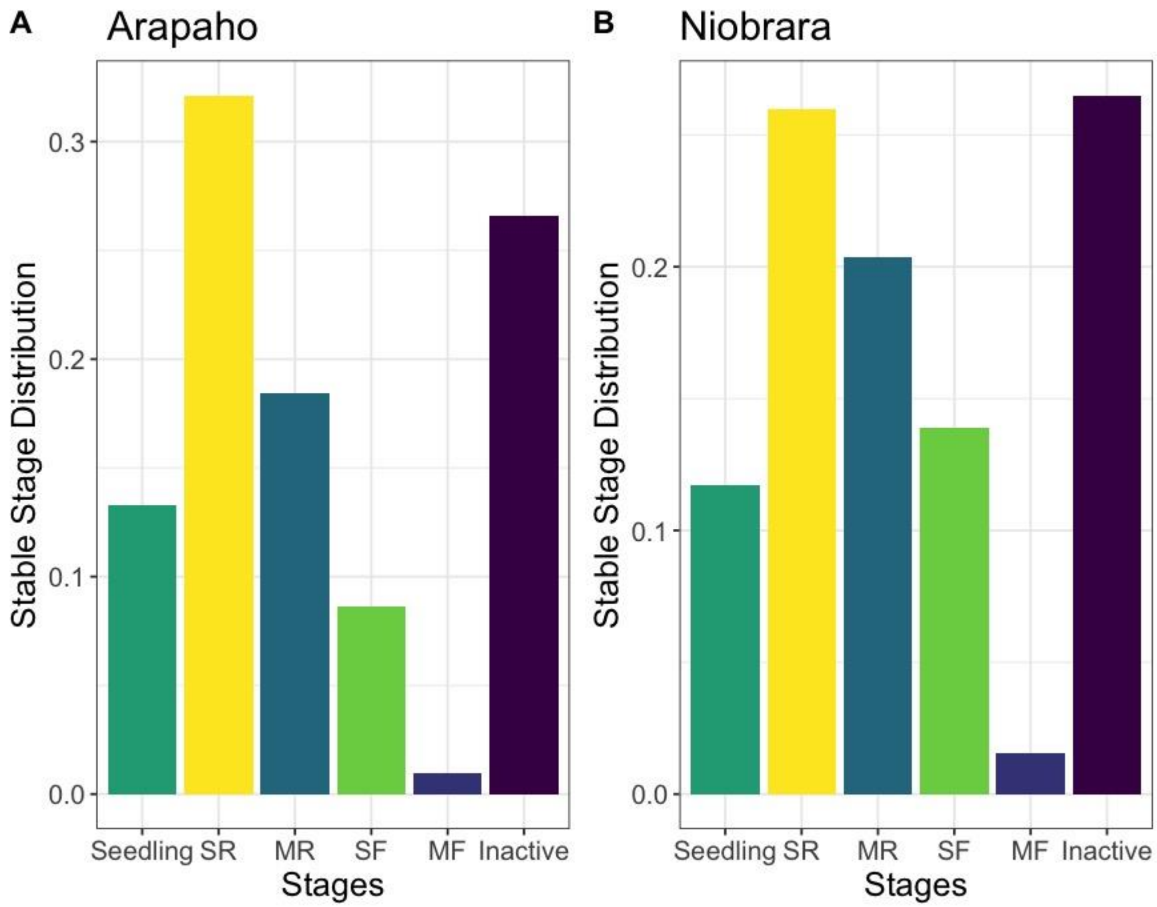


FIGURE S5: The proportion of individuals expected in each stage at a stable stage distribution.
